# History modulates early sensory processing of salient distractors

**DOI:** 10.1101/2020.09.30.321729

**Authors:** Kirsten C.S. Adam, John T. Serences

## Abstract

To find important objects, we must focus on our goals, ignore distractions, and take our changing environment into account. This is formalized in models of visual search whereby goal-driven, stimulus-driven and history-driven factors are integrated into a priority map that guides attention. Stimulus history robustly influences where attention is allocated even when the physical stimulus is the same: when a salient distractor is repeated over time, it captures attention less effectively. A key open question is how we come to ignore salient distractors when they are repeated. Goal-driven accounts propose that we use an active, expectation-driven mechanism to attenuate the distractor signal (e.g., predictive coding), whereas stimulus-driven accounts propose that the distractor signal is attenuated due to passive changes to neural activity and inter-item competition (e.g., adaptation). To test these competing accounts, we measured item-specific fMRI responses in human visual cortex during a visual search task where trial history was manipulated (colors unpredictably switched or were repeated). Consistent with a stimulus-driven account of history-based distractor suppression, we found that repeated singleton distractors were suppressed starting in V1, and distractor suppression did not increase in later visual areas. In contrast, we observed signatures of goal-driven target enhancement that were absent in V1, increased across visual areas, and were not modulated by stimulus history. Our data suggest that stimulus history does not alter goal-driven expectations, but rather modulates canonically stimulus-driven sensory responses to contribute to a temporally-integrated representation of priority.

**Significance Statement:** Visual search refers to our ability to find what we are looking for in a cluttered visual world (e.g., finding your keys). To perform visual search, we must integrate information about our goals (e.g., ‘find the red key-chain’), the environment (e.g., salient items capture your attention), and changes to the environment (i.e., stimulus history). Although stimulus history impacts behavior, the neural mechanisms that mediate history-driven effects remain debated. Here, we leveraged fMRI and multivariate analysis techniques to measure history-driven changes to the neural representation of items during visual search. We found that stimulus history influenced the representation of a salient ‘pop-out’ distractor starting in V1, suggesting that stimulus history operates via modulations in early sensory processing rather than goal-driven expectations.

## Introduction

At any moment we can selectively attend only a small fraction of available perceptual inputs, so we need to select a subset of information and discard irrelevant information. This capacity limit poses a significant computational challenge, particularly given that perceptual inputs constantly change as we move through the world. Given our constantly changing surroundings, one particularly useful computational strategy is to discard information that stays the same over time. For example, when searching for sea glass at the beach, irrelevant but salient information (e.g., a red plastic bottle-cap) may grab our attention. But, if we repeatedly encounter the same irrelevant information (e.g., the beach is littered with red bottle-caps), then we can come to ignore initially salient distractors. Extensive evidence suggests that stimulus history robustly modulates behavior: An initially salient color distractor no longer captures our attention after we have seen it many times (Geyer et al., 2006; Vatterott and Vecera, 2012; Geng, 2014; Gaspelin et al., 2015, 2017; Wang and Theeuwes, 2018a; Failing et al., 2019a; Geng et al., 2019; Van Moorselaar and Slagter, 2020). Yet, debate persists as to how this history-driven behavioral modulation is achieved: Do we use an active, expectation-based mechanism to suppress the salient distractor signal when it is repeated (e.g., predictive coding), or, is the distractor signal passively attenuated because of changes to neural activity with repetition (e.g., adaptation)? Here, we leverage item-specific, neural estimates of priority, to test competing hypotheses of how stimulus history alters attentional priority.

Models of visual search hypothesize that we integrate information about what is relevant *(goal-driven* or *‘top-down’ factors*), what is salient given local image statistics *(stimulus-driven* or *‘bottom-up’ factors*), and what has occurred in the past (*history-driven factors*) via an integrated, topographically organized “priority map” (Treisman and Gelade, 1980; Wolfe, 1994; Itti and Koch, 2000; Fecteau and Munoz, 2006; Serences and Yantis, 2006; Awh et al., 2012). Note, some work uses the terms ‘saliency’ and ‘priority’ interchangeably, whereas other work uses these terms to refer to distinct concepts. Here, we use ‘priority’ to refer to the integration of goal-driven and stimulus-driven task factors, and ‘saliency’ to refer to strictly image-computable, stimulus-driven task factors (Serences and Yantis, 2006). Although both stimulus-driven and goal-driven information is represented to some extent in many cortical regions (Silver et al., 2005; Serences and Yantis, 2006, 2007; Saproo and Serences, 2010; Bogler et al., 2011; Sprague and Serences, 2013; Sprague et al., 2018b), areas of parietal cortex (e.g., LIP, IPS) are hypothesized to be ideal candidates for integrating information about stimulus-driven sensory inputs from occipital cortex and information about goals from pre-frontal cortex (Ipata et al., 2006, 2009; Bisley and Mirpour, 2019; Theeuwes, 2019).

History-driven effects have only recently been added to models of visual search, in part because these effects do not wholly fit within a ‘goal-driven’ versus ‘stimulus-driven’ dichotomous framework (Awh et al., 2012; Geng, 2014; Le Pelley et al., 2016; Geng et al., 2019; Van Moorselaar and Slagter, 2020). Rather, history-driven effects rely on both current sensory input and prior experiences. Thus, debate persists about whether stimulus history influences attentional priority by co-opting elements of stimulus-driven computations, goal-driven computations, or another pathway altogether (Gaspelin et al., 2015; Gaspelin and Luck, 2018; Wang and Theeuwes, 2018b; Geng et al., 2019; Theeuwes, 2019; Van Moorselaar and Slagter, 2020).

Goal-driven accounts propose that we exploit voluntary selection mechanisms to incorporate information about history-driven task factors into priority maps. Earlier work has shown how voluntary attention may be used to enhance the target item relative to the other distractor items: when looking for a particular target, one may form a “template” of that feature and use this template to voluntarily up-regulate relevant portions of the visual field (Pashler and Shiu, 1999; Downing, 2000; Soto et al., 2005; Olivers et al., 2006; Carlisle et al., 2011; Beck et al., 2012). Similar voluntary selection mechanisms might likewise be used to suppress a distractor signal when it is repeated, either directly or indirectly. First, distractor suppression could arise *indirectly* because of predictive coding and biased competition (Desimone and Duncan, 1995; Spratling, 2008; Summerfield and de Lange, 2014). In this account of history-driven distractor suppression, participants could use their expectations about the upcoming, repeated stimulus futures to more strongly enhance the target and, consequently, the competing distractor would be automatically suppressed due to inter-item competition (Spratling, 2008). Second, distractor suppression could arise *directly*, via a top-down suppression signal for a specific feature. This direct suppression signal is sometimes referred to as a “negative search template” (Arita et al., 2012; Moher and Egeth, 2012; Reeder et al., 2017; Conci et al., 2019, but see: Beck and Hollingworth, 2015; Becker et al., 2015). Critically, in either the direct or indirect case, we would expect to observe a similar neural signature at the level of population codes measured with fMRI for both of these goal-driven accounts. Specifically, we should observe that distractor suppression effects are greater in later visual areas (e.g., IPS0) than in earlier visual areas (e.g., V1), consistent with a goal-driven signal (e.g., Silver et al., 2005; Sprague et al., 2018b). In the case of the predictive coding / biased competition model (Spratling, 2008), we would further predict that target enhancement and distractor suppression should be yoked, whereby greater distractor suppression will be accompanied by greater target enhancement as arrays are repeated.

In contrast, stimulus-driven accounts instead suggest that history-driven distractor suppression can arise from passive changes to neural activity as stimuli are presented over time (Turatto and Pascucci, 2016; Turatto et al., 2018; Wang and Theeuwes, 2018a; Failing et al., 2019a; Won and Geng, 2020). For example, some work suggests that even passive, task-irrelevant exposure to a particular feature may be sufficient to alter attentional guidance and search behaviors (Engel and Furmanski, 2001; Grill-Spector and Malach, 2001; Gardner et al., 2005; Kristjansson et al., 2007; Turatto and Pascucci, 2016; Turatto et al., 2018; Won and Geng, 2020). Although the effects of adaptation for single stimuli are well understood, how adaptation affects saliency in multi-item displays has only recently been considered. Yet, emerging evidence suggests that altered firing rates from simple sensory adaptation effects could alter inter-item competition (Solomon and Kohn, 2014) which, in turn, could alter stimulus saliency and behavior (Treisman and Gelade, 1980; Wolfe, 1994; Itti and Koch, 2000; Li, 2002; Zhang et al., 2012).

Here, we tackle the debate about history-driven distractor suppression from a new angle: we measured neural activity via fMRI in human subjects performing a visual search task in order to estimate item-specific changes to neural priority across the visual stream. Critically, we manipulated trial history so that we could compare neural responses to physically identical displays (e.g., green target, red singleton distractor) as a function of trial history (i.e., whether the colors of preceding displays repeated or varied).

## Materials and Methods

### Participants

#### Experiment 1a: MRI experiment

Healthy volunteers (*n* = 12; 9 female; mean age = 25.3 years [SD = 2.5, min = 21, max = 30]; all right-handed; normal or corrected-to-normal visual acuity; normal color vision) participated in three ∼2 hour sessions at the Keck Center for fMRI on the University of California San Diego (UCSD) campus, and were compensated $20/hr. Procedures were approved by the UCSD Institutional Review Board, and participants provided written informed consent. Sample size was determined by a power analysis on data from Sprague et al.(Sprague et al., 2018b) where achieved power (1-ß) to detect a within-subjects attention modulation using an inverted encoding model was 83% (across 10 ROIs) with n=8. We planned for n =11 to achieve estimated 90% power (rounded up to n = 12 to satisfy our counter-balancing criteria).

#### Experiment 1b: Behavior only

Healthy volunteers (*n* = 24; 21 female; mean age = 19.8 years [SD = 1.5, min = 18, max = 24]; normal or corrected-to-normal visual acuity; normal color vision; handedness not recorded) participated in one 1.5-hour experimental session in the Department of Psychology on the UCSD campus, and were compensated with course credit. There were no duplicate participants across experiments. Procedures were approved by the UCSD IRB, and all participants provided written informed consent. A sample size of 24 was chosen *a priori* based on published papers(Gaspelin et al., 2015).

### Session procedures

#### Exp 1a, Retinotopy session

Participants completed one retinotopic mapping session prior to participation in the experimental sessions, following standard procedures(Engel et al., 1994; Swisher et al., 2007). Some participants had already completed a retinotopy session as part of prior studies in the lab; this session was used if available. Retinotopy data were used to identify retinotopic ROIs (V1-V3, V3AB, hV4, VO1, VO2, LO1, LO2, TO1, TO2, IPS0-4). During each session, participants viewed flickering checkerboards. On meridian mapping runs, a “bowtie” checkerboard alternated between the horizontal and vertical meridians. On polar angle mapping runs, a checkerboard wedge slowly rotated in a clockwise or counterclockwise direction. On eccentricity mapping runs, a “donut” checkerboard began near fixation and its radius slowly expanded outward. A high-resolution anatomical scan was collected for functional alignment. Anatomical and functional retinotopy analyses were performed using custom code calling existing FreeSurfer and FSL functions. Functional retinotopy data were used to draw ROIs, but only voxels that were also visually responsive to experimental localizers (below) were analyzed further.

#### Exp 1a, Main MRI session

Participants completed two experimental sessions. In each session, they completed 2 runs of the item position localizer, 4 runs of the spatial location localizer, and 8 runs of the search task (4 runs “color variable”, 4 runs “color constant”). When time allowed, extra localizer runs were collected. Some participants also took part in an unrelated study in which additional localizers were collected. Across the two sessions, participants completed 16 runs of visual search (M = 1,1152 trials, SD = 0), an average of 11.2 runs of the spatial location localizer (M = 1,072 trials, SD = 298, min = 768, max = 1,536), and an average of 4.3 item position localizer runs (M = 381 trials, SD = 43, min = 352, max = 440).

#### Exp 1b

Participants completed 12 blocks of the search task (6 blocks “color variable”, 6 blocks “color constant”).

### Stimuli and task procedures

#### Experiment 1a: MRI

Stimuli were projected on a 21.5 × 16 cm screen mounted inside the scanner bore. The screen was viewed from a distance of ∼47 cm through a mirror. Stimuli were generated in MATLAB (2017b, The MathWorks, Natick, MA) with the Psychophysics toolbox (Brainard, 1997; Pelli, 1997) on a laptop running Ubuntu. Responses were collected with a 4-button button box. Stimuli for each task are shown in Figure 1.

**Figure 1.**
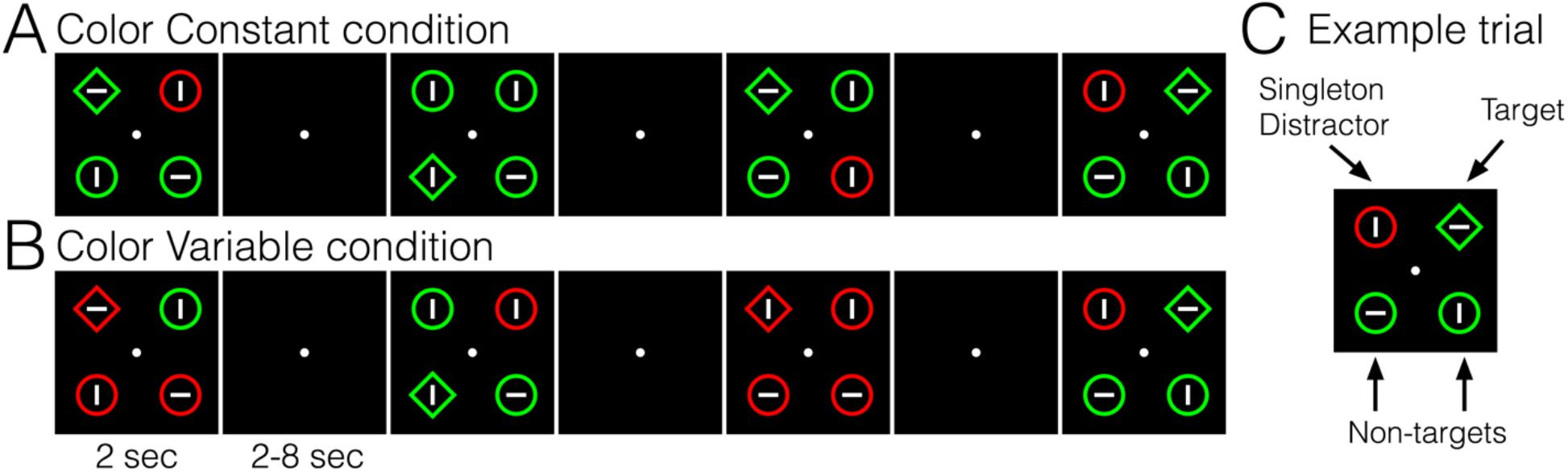
Visual search task stimuli. On each trial, participants viewed a 4-item array and reported the orientation of the line inside the diamond-shaped target (horizontal or vertical). **(A)** In the color constant condition, colors of targets and singleton distractors were fixed throughout the run. **(B)** In the color variable condition, colors of targets and singleton distractors swapped randomly from trial to trial. **(C)** An example trial with labels for the target, singleton distractor, and non-target items.

#### Item position localizer

Participants viewed reversing checkerboards (4 Hz flicker) which occupied the locations of the items in the search task (each item radius = 2.5° placed on an imaginary circle 7° from fixation, with one item in each of the 4 quadrants on the circle). Participants were shown items on 2 alternating diagonals (i.e., items in Quadrants 1 and 3 and then Quadrants 2 and 4) for 3 sec each. There were 88 stimulus presentations within each run. Participants were instructed to attend to both items, and to press a button if either item briefly dimmed. A brief (250 ms) dimming occurred on 1 of the 2 items for 25% of stimulus presentations.

#### Spatial location localizer

Participants viewed a reversing checkerboard wedge (flicker = 4 Hz; white & black checkerboards) at one of 24 positions. Checkerboard positions were equally spaced along a circle with radius = 7°, and wedges were non-overlapping (i.e., each wedge’s width along the circle filled a 15° arc and was ∼5° of visual angle in height). The wedge stayed at one position for 3 sec, then moved to a different position (with the constraint that back-to-back positions must be in different quadrants). There were 96 wedge presentations within each run. Participants were instructed to attend to the fixation point; if the fixation point’s color changed (increase or decrease in brightness), they pressed a button on the button box. A total of 20 fixation point color changes occurred throughout each run; changes to the fixation cross happened at random times with respect to wedge stimulus onsets. We chose to have participants attend fixation, rather than the stimulus position, during the localizer task to reduce contamination of eye movements on any observed decoding effects (Mostert et al., 2018). Generally, systematic eye movement biases are absent or greatly attenuated when participants attend fixation and ignore the peripheral stimulus. With this cross-task training and testing scheme, we would expect that decoding should impaired for all 4 items if participants moved their eyes in the visual search task. Thus, this training-testing scheme protects against the possibility that item-specific target enhancement or distractor suppression effects could be driven by eye movements to a particular item in the display.

#### Search task

Participants performed a variant of the additional singleton search task (Theeuwes, 1992). On each trial, participants saw a search array containing 4 items (item colors were red, RGB = 255,0,0, or green, RGB = 0,255,0, and presented on a black background, RGB = 0,0,0). The items (2.4° radius) were placed on an imaginary circle 7° from fixation with 1 item in each visual quadrant (i.e., 45°, 135°, 225° & 315°). Participants fixated a small, gray dot (.2°) throughout each run. Participants searched for a “target” (the diamond-shaped item) among distractor items and reported the orientation of the small line inside (line size = .08° x .94°; orientation = horizontal or vertical) by pressing one of two buttons. Non-singleton distractors, ‘non-targets’, had the same color as the shape-defined target (e.g., green circles). A “singleton distractor” was present on 66.67% of trials, and was a color singleton (e.g., red circle). Stimuli are illustrated in Figure 1. Note, throughout the manuscript, we will use the word “distractor” to refer to the color-singleton distractors, whereas non-singleton distractors will be referred to simply as “non-targets”. Target location (quadrant 1-4), distractor location relative to the target (−90°, +90°, or +180°), distractor presence (66.67% present), and the orientation of the line inside the target (horizontal or vertical) were fully counterbalanced within each run, for a total of 72 trials per run. Search set size was held constant at 4 items. The search array was presented for 2 sec followed by a blank inter-trial interval (equal probability of 2, 3.2, 5, or 8 sec).

We manipulated the degree to which participants were behaviorally captured by the distractor by changing trial history. In “color variable” runs, the colors of targets and distractors swapped unpredictably. In “color constant” runs, the colors of targets and distractors were fixed throughout the run (e.g., the targets and non-singleton distractors were always green and the singleton distractor was always red). Based on prior work (Vatterott and Vecera, 2012; Gaspelin et al., 2017), we expected to observe robust behavioral capture by the singleton distractor in the color variable runs and no behavioral capture in the color constant runs.

Run types were blocked and partially counterbalanced within and across sessions, such that the order of the 2 conditions would be balanced across the 2 sessions for each participant. For example, if in Session 1 a participant first received 4 color variable runs followed by 4 color constant runs (red), then in Session 2 they would first receive 4 color constant runs (green) followed by 4 color variable runs.

#### Experiment 1b: Behavior

Participants performed the same additional singleton search task described above. Participants viewed the stimuli on CRT monitors (39 × 29.5 cm) from a distance of ∼52 cm. Stimulus parameters (size, color) and trial timing were matched to the fMRI experiment. Each experimental block contained a total of 48 search trials. Participants performed a total of 12 blocks of trials (6 color variable, 3 color constant with red targets, 3 color constant with green targets). The color constant and color variable conditions were blocked and counterbalanced across participants (half of participants received the color variable condition first).

### Magnetic resonance imaging acquisition parameters

Scans were performed on a General Electric Discovery MR750 3.0T scanner at the Keck Center for Functional Magnetic Resonance Imaging on the UCSD campus. High-resolution (1mm^3^ isotropic) anatomical images were collected as part of the retinotopy session. Most participants’ (10 of 12) anatomical images were collected with an Invivo 8-channel head coil; 2 participants’ anatomical images were collected with a Nova Medical 32-channel head coil (NMSC075-32-3GE-MR750). GE’s “Phased array Uniformity Enhancement” (PURE) method was applied to anatomical data acquired using the 32-channel coil in an attempt to correct inhomogeneities in the signal intensity. Functional echo-planar imaging (EPI) data were collected with the Nova 32 channel coil using the GE multiband EPI sequence, using nine axial slices per band and a multiband factor of eight (total slices = 72; 2 mm^3^ isotropic; 0 mm gap; matrix = 104 × 104; field of view = 20.8 cm; repetition time/echo time (TR/ TE) = 800/35 ms, flip angle = 52°; in-plane acceleration = 1). The initial 16 TRs in each run served as reference images for the transformation from *k*-space to image space. Un-aliasing and image reconstruction procedures were performed on local servers and on Amazon Web Service servers using code adapted from the Stanford Center for Cognitive and Neurobiological Imaging (CNI). Forward and reverse phase-encoding directions were used during the acquisition of two short (17 sec) “top-up” datasets. From these images, susceptibility-induced off-resonance fields were estimated (Andersson et al., 2003) and used to correct signal distortion inherent in EPI sequences, using FSL top-up (Smith et al., 2004; Jenkinson et al., 2012).

### Pre-processing

Pre-processing of imaging data closely followed published lab procedures (Rademaker et al., 2019) using FreeSurfer and FSL. We performed cortical surface gray-white matter volumetric segmentation of the high-resolution anatomical volume from the retinotopy session using FreeSurfer’s “recon-all” procedures (Dale et al., 1999). The first volume of the first functional run from each scanning session was coregistered to this common T1-weighted anatomical image. To align data from all sessions to the same functional space, we created transformation matrices with FreeSurfer’s registration tools (Greve and Fischl, 2009), and used these matrices to transform each four-dimensional functional volume using FSL’s FLIRT (Jenkinson and Smith, 2001; Jenkinson et al., 2002). After cross-session alignment, motion correction was performed using FSL’s McFLIRT (no spatial smoothing, 12 degrees of freedom). Voxelwise signal time-series were normalized via Z-scoring on a run-by-run basis. Analyses after preprocessing were performed using custom scripts in MATLAB 2018A.

### fMRI analyses: Inverted encoding model

#### Voxel selection for Decoding ROIs

We defined visual ROI’s using data from the retinotopy session following published lab procedures (Sprague and Serences, 2013; Rademaker et al., 2019). From these retinotopically-derived ROI’s, we chose the subset of voxels that were spatially selective for the stimuli used in this task. We thresholded voxels using the independent mapping task data. We ran a one-way ANOVA with factor Quadrant on each voxel; significant voxels (*p* < .05 uncorrected) were retained for analysis. For the aggregate analyses, we *a priori* created an early visual cortex ROI (all spatially selective voxels from V1-V3) and a parietal cortex ROI (all spatially selective voxels from IPS0-3). For individual ROI analyses, we used all individual retinotopic ROIs for which there were a minimum of 90 spatially selective voxels per participant: V1, V2, V3, V3AB, hV4, and IPS0.

#### Inverted Encoding Model

Following prior work (Brouwer and Heeger, 2009; Sprague and Serences, 2013), we used an inverted encoding model to estimate spatially-selective tuning functions from multivariate, voxel-wise activity within each ROI. We assumed that each voxel’s activity reflects the weighted sum of 24 spatially selective channels, each tuned for a different angular location. These information channels are assumed to reflect the activity of underlying neuronal populations tuned to each location.

We modeled the response profile of each spatial channel as a half sinusoid raised to the 24^th^ power:

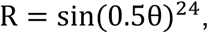

where θ is angular location (0–359°, centered on each of the 24 bins from the mapping task), and *R* is the response of the spatial channel in arbitrary units.

Independent training data *B*_*1*_ were used to estimate weights that approximate the relative contribution of the 24 spatial channels to the observed response at each voxel. Let *B*_*1*_ (*m* voxels × *n*_*1*_ observations) be the activity at each voxel for each measurement in the training set, *C*_*1*_ (*k* channels × *n*_*1*_ observations) be the predicted response of each spatial channel (determined by the basis functions) for each measurement, and *W* (*m* voxels × *k* channels) be a weight matrix that characterizes a linear mapping from “channel space” to “voxel space”. The relationship between *B*_*1*_, *C*_*1*_, and *W* can be described by a general linear model:

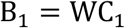

We obtained the weight matrix through least-squares estimation:

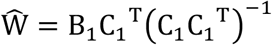

In the test stage, we inverted the model to transform the observed test data *B*_*2*_ (*m* voxels × *n*_*2*_ observations) into estimated channel responses, *C*_*2*_ (*k* channels × *n*_*2*_ observations), using the estimated weight matrix, Ŵ, that we obtained in the training phase:

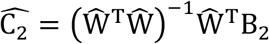

Each estimated channel response function was then circularly shifted to a common center by aligning the estimated channel responses to the channel tuned for target location.

#### Model training and testing

We trained the IEM using independent mapping task data and tested the model using single trial search-task data (average of 4 to 10 TR’s after search array onset). We then shifted and averaged the search task data so that like trials were aligned (e.g., rotate and average all trials with target-distractor distance of 90). To reduce idiosyncrasies of only having 1 test set, we iterated the analysis by leaving out 1 block of training data and 1 block of test data, looping through all possible combinations (e.g., for each 1 block of left out training data, we left out each possible block of test data on different runs of the loop).

## Results

### Behavior

Subjects performed a variant of the additional singleton search task (Theeuwes, 1992) (Figure 1A) in which they searched for a target (diamond) among non-targets (circles). On each trial, the participant reported via button-press the orientation of the line inside the diamond target (vertical or horizontal). On 66.67% of trials, one of the non-targets was uniquely colored (“singleton distractor present”, e.g., one red distractor, two green non-targets, and one green target item). Behavioral capture was quantified as slowed response times (RTs) when the distractor was present versus absent. In addition to examining the basic capture effect, a key goal of this work was to examine modulation of capture by trial history (Vatterott and Vecera, 2012; Gaspelin et al., 2015, 2017). Prior work has shown that participants can learn to suppress a distractor (i.e., no RT difference for singleton distractor present versus absent trials) when the same distractor color or distractor location is repeated over many trials (Vatterott and Vecera, 2012; Gaspelin et al., 2015, 2017). Building on this work, we included two key task conditions in a counterbalanced, block-wise fashion to manipulate trial history and behavioral capture while using identical stimulus arrays (e.g., green target, red distractor). In the color constant condition (Figure 1A), the array colors stayed constant throughout the block (e.g., green target, green non-target items, red distractor). In the color variable condition (Figure 1B), the array colors randomly varied from trial to trial. Based on prior work, we expected robust capture in the color variable condition, and little or no capture in the color constant condition (Vatterott and Vecera, 2012; Gaspelin et al., 2015, 2017).

Replicating prior work, we found significant behavioral capture that was modulated by trial history (Geyer et al., 2006; Vatterott and Vecera, 2012; Goschy et al., 2014; Gaspelin et al., 2015, 2017; Wang and Theeuwes, 2018a; Failing et al., 2019a). In our MRI sample (Exp 1a, Figure 2A & 2C), we observed significant behavioral capture in the color variable condition, with longer RT’s for distractor present versus distractor absent trials (*M* = 32.8 ms, *SD* = 25.5 ms, *p* = .001, *d* = 1.28), but capture was not significant in the color constant condition (*M* = 10.8 ms, *SD* = 18.5 ms, *p* = .07, *d* = .59). Importantly, capture was significantly larger for color variable vs. color constant runs (*p* = .009, *d =*.91). We replicated this pattern of findings in the behavior-only experiment (Exp. 1b, Figure 2B & 2D), with robust capture for ‘color variable’ (*p* < 1 × 10^−5^, *d* = 1.31), no significant capture for ‘color constant’ (*p* = .1, *d* = .32), and larger capture for color variable vs. constant (*p* = .002, *d* = .71). Participants in both experiments were accurate overall (>90%), and there was no evidence of a speed-accuracy trade-off (Figure 2-1).

**Figure 2.**
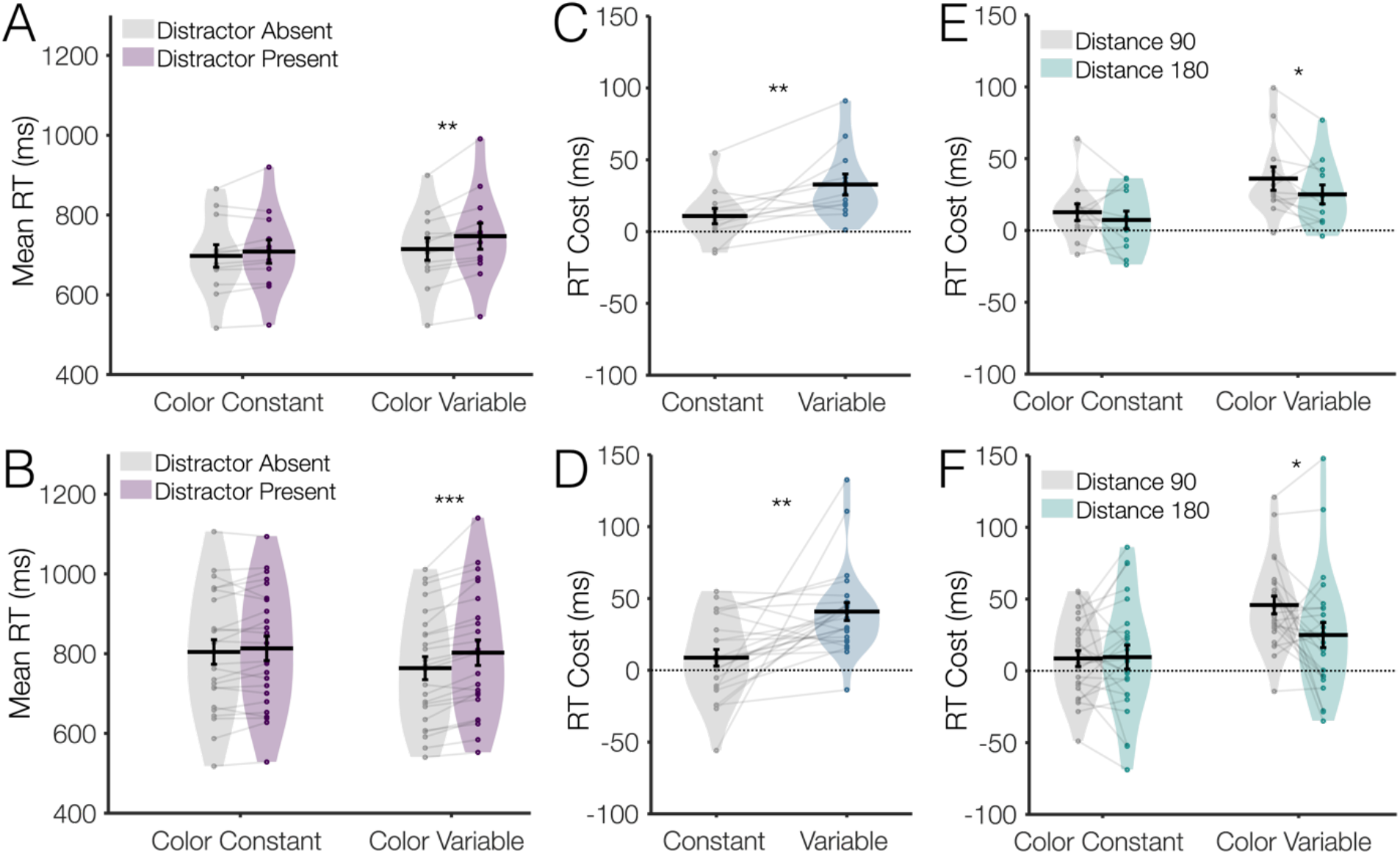
Behavioral capture during the visual search task. **(A)** In the main MRI Experiment (Exp 1a), participants were significantly captured by the salient singleton distractor in the color variable condition, but not in the color constant condition. **(B)** This pattern replicated in the behavior-only experiment (Exp 1b). **(C-D)** Capture costs (RT Difference for distractor present – absent trials) were significantly larger in the color variable than in the color constant condition in Exp 1a **(C)** and Exp 1b **(D). (E-F)** Capture costs (RT Difference for distractor present – absent trials) were significantly modulated by the distance between the target and distractor in the color variable condition both in Exp 1a **(E)** and Exp 1b **(F)**. Violin plot shading shows range and distribution of the data; dots represent single subjects; black error bars indicate ±1 SEM. Asterisks depict significance for uncorrected post-hoc comparisons between adjacent bars within each experiment, * *p* < .05, ** *p* < .01, *** *p* < .001.

In addition to the key modulation of capture as a function of stimulus history, we also replicated prior findings that the degree of capture is significantly modulated by the physical distance between the target and the distractor (Mounts, 2000; Turatto and Galfano, 2001; Wang and Theeuwes, 2018a; Failing et al., 2019b), with larger capture for distractors nearer the target (Figure 2E-F). We ran a repeated measures ANOVA including both experiments (n=36). Including Experiment as a factor revealed no experiment main effects or interactions (p > .2), so the two experiments were combined for further analyses of the behavioral data (although Figure 2 shows data from the two experiments separately). There was a significant effect of Condition (larger capture for color variable than color constant), *p <* 1×10^−4^, a main effect of Distance (larger capture for 90° than 180°), *p* = .037, η^2^_p_ = .12, and an interaction between Condition and Distance (greater distance effect in the color variable condition), *p* = .014, η^2^_p_ = .16.

### fMRI results: Model estimates of spatial position in the independent mapping task

We opted for a multivariate model-based approach to estimate the amount of information encoded in voxel activation patterns about each of the 4 stimuli in the search array, as such multivariate approaches are more sensitive than just computing the univariate mean response across all voxels (Cox and Savoy, 2003; Haynes and Rees, 2005; Kamitani and Tong, 2005; Norman et al., 2006; Serences and Saproo, 2012; Tong and Pratte, 2012). For example, item-specific information has been observed using multivariate methods even in the absence of univariate changes (Lewis-Peacock and Postle, 2012; Emrich et al., 2013) (but for univariate analyses of the present data, see extended data Figure 3-1). We opted for an inverted encoding model (IEM) approach (Sprague et al., 2018a, 2019), as opposed to Bayesian or other decoders (van Bergen et al., 2015; van Bergen and Jehee, 2019), because this approach allowed us to easily derive a separate estimate of the information encoded about each of the 4 simultaneously presented items from the search array in the main analysis (Sprague et al., 2019). For further discussion of IEM model assumptions and best practices, see: (Sprague et al., 2018a, 2019).

**Figure 3.**
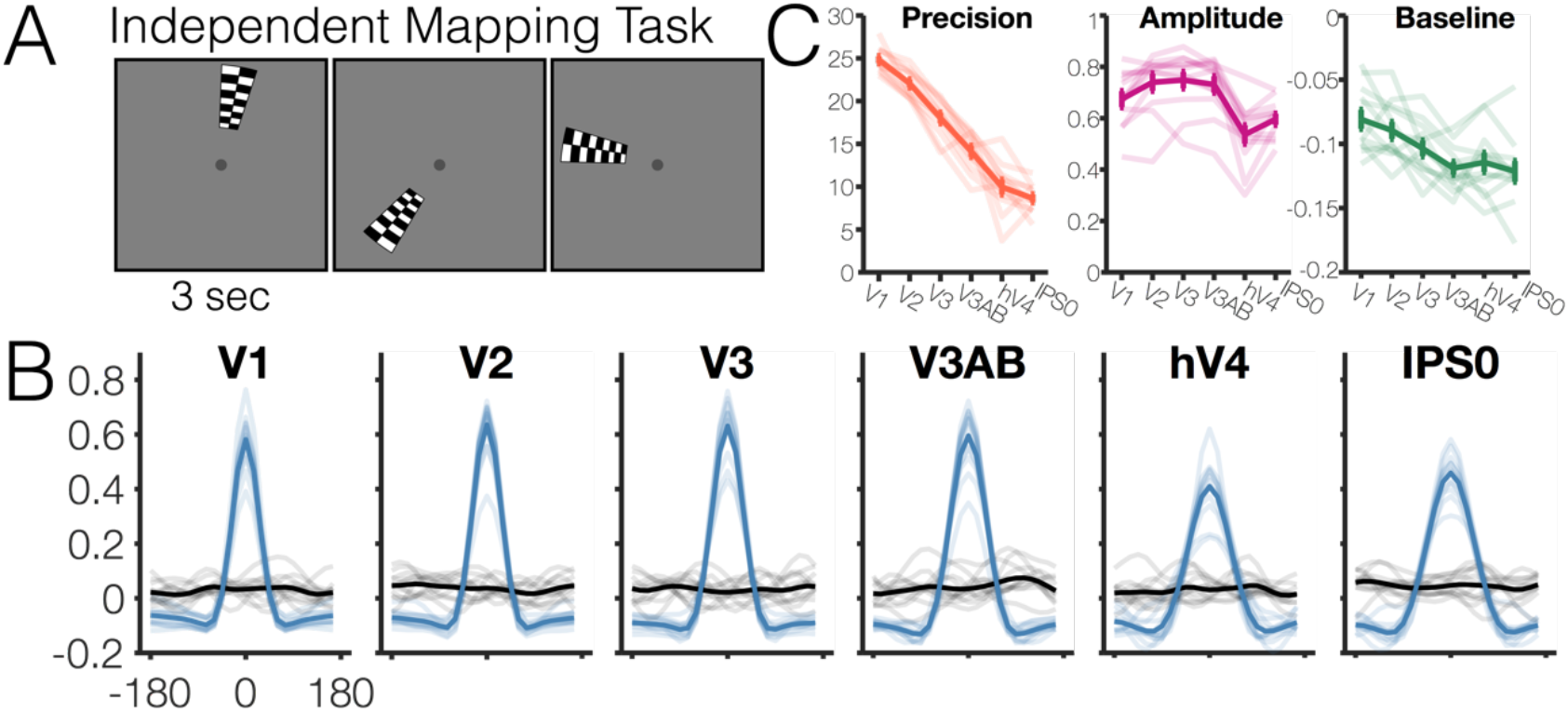
Single-item model estimates training and testing within the independent mapping task. **(A)** Independent mapping task used to train the model to estimate spatial position of 4 search array items. Participants viewed a flickering checkerboard which could appear at one of 24 positions around an imaginary circle. **(B)** Blue lines: Model estimates of viewed spatial position training and testing within the independent mapping task. Single-trial model estimates for each subject are aligned to 0 degrees and averaged. Black lines: Model estimates for shuffled training labels. Opaque lines = group average; semi-transparent lines = individual subjects. **(C)** Descriptive statistics for best fit von Mises parameters (precision [κ], amplitude, baseline) to model estimates in panel B. Error bars indicate ±1 SEM; the opaque line shows the group average; semi-transparent lines show individual subjects.

In our key analyses of the fMRI data, we used an independent mapping task to train a model of spatial position from which we estimated the relative priority of all item positions within the visual search array. During the independent mapping task, observers viewed a flickering checkerboard wedge that was presented at 1 of 24 positions on an imaginary circle around fixation (Figure 3A). We first checked that we observed robust estimates of spatial position when training and testing within the independent mapping task (leave 1 run out, see section ‘Inverted Encoding Model’). We observed robust model-based estimates of spatial position for all ROIs (Figure 3B). Parameters from the best-fitting von Mises distribution to each region-of-interest (ROI) are depicted in Figure 3C (model fits, linear classifier results, and von Mises parameters for shuffled data are shown in extended data Figure 3-2). There was an effect of ROI on precision such that spatial position was represented less precisely in later visual areas (*p* < 1×10^−5^, where precision is the concentration parameter κ of the best fitting von Mises, with higher values indicating a more precise function). There was also an effect of ROI on the amplitude and baseline measures of the model-based estimates of spatial position (*p* < 1×10^−5^), and all 3 parameters significantly differed from zero across all ROIs (p < 1×10^−5^). These results, particularly the observation of amplitudes greater than 0, confirmed that activation patterns in all examined regions encode information about spatial position.

Unlike the single item model estimates that were derived based on the independent mapping task (Figure 3), we could not fit a simple, uni-modal Gaussian function to model-based estimates derived from the search task data because 4 peaks in the model output were expected – one for each item in the search array. As such, we first conducted simulations to ensure that we would be able to measure putative changes to individual item representations (e.g. target enhancement, distractor suppression), despite multiple item representations contributing to the aggregate 4-item model estimates. To do so, we used data from the independent mapping task to generate predictions for observed model responses in a 4-item array. For each ROI, we took the 1-item model response derived from the independent mapping task, replicated this model response four times (once at each of the four search array positions), and took the average of all 4 shifted 1-item model response lines to generate a single 4-item model prediction. In addition, we systematically varied the strength of the simulated response to each item to ensure that we were able to recover a corresponding change in the item-specific responses estimated from the aggregate 4-item model estimate (Figure 4; extended data Figure 4-1).

**Figure 4.**
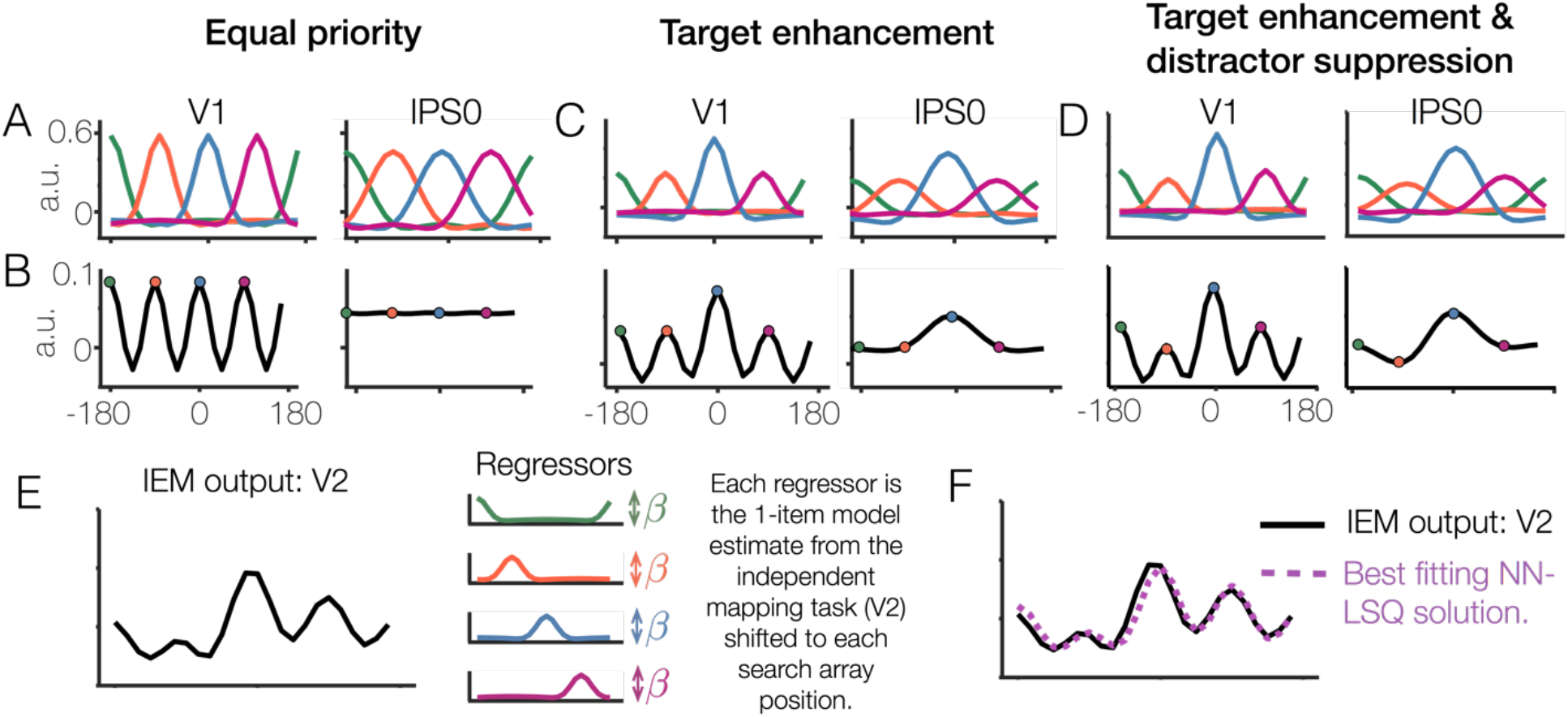
Generating predictions for 4-item model estimates by averaging single-item model estimates from the independent mapping task. **(A)** Average from the independent mapping task plotted at 4 hypothetical item locations. Here, these 4 “items” are represented with equal priority. (B) Hypothetical observed response when measuring a single trial containing the 4 items presented simultaneously. This line is the average of all lines in Panel A. (C) The same as panels A and B, but with the item at position 0 assigned a higher response amplitude than the other three items. (D) The same panels as A and B, but with both an enhanced item at position 0 and a suppressed item at position -90. (E) Actual IEM model output for 4-item search arrays in V2 (Target plotted at 0, distractor plotted at -90). To estimate the strength of each of the 4 underlying item representations, one can simply measure the height (a.u.) at expected item peaks (i.e., - 180, -90, 0, and 90). Alternatively, one may use a non-negative least squares solution to estimate weights for a regressor for each of the 4 item positions. Each regressor is the 1-item IEM output from the independent mapping task within the same region (e.g., V2), shifted to the appropriate item location. (F) Example IEM output and best-fitting non-negative least squares solution with 4 item regressors.

These simulations revealed clearly separable peaks for all four items in early areas like V1, where spatial precision is high (Figure 4A-B, left panel). In contrast, identifying clear peaks in later areas like IPS0 was difficult when the response to all items was equivalent (Figure 4A-B, right panel). However, if one item evoked a larger or smaller response than the other items, as would be expected with target enhancement or distractor suppression, then clear and measurable changes to the aggregate 4-item model estimates emerged (Figure 4C). Further simulations showed that we could detect smaller changes to one item (e.g., distractor suppression) in the presence of larger changes to another item (e.g., target) by measuring the response amplitude at each expected item’s peak. In V1, this is clearly seen in the peak response to each item; in later areas such as IPS0, such changes manifest as a large central peak that is skewed by the neighboring items’ smaller changes (Figure 4D).

We also used a general linear model (GLM) to estimate best-fitting gain factors for each of the 4 hypothesized item representations by fitting an aggregate function and allowing one parameter in the GLM to scale the response associated with each item. This is essentially the inverse of the simulations described above: For a given aggregate response (i.e., the response of each of the 24 spatial channels when shown a given 4-item search array), we used a non-negative least squares solution(Lawson and Hanson, 1974) to estimate the contribution of each of the 4 item positions (calculated from the 1-item localizer task) to the observed 4-item search array response (Figure 4E). This analysis yielded similar results to the simple approach of comparing the height at each expected item peak (extended data Figure 5-4). Thus, using either the raw amplitude at expected peaks or a GLM-based approach, we determined that we should be able to accurately characterize situations in which there was no modulation of target and distractor responses as well as situations in which there was a significant modulation of target and/or distractor responses.

**Figure 5.**
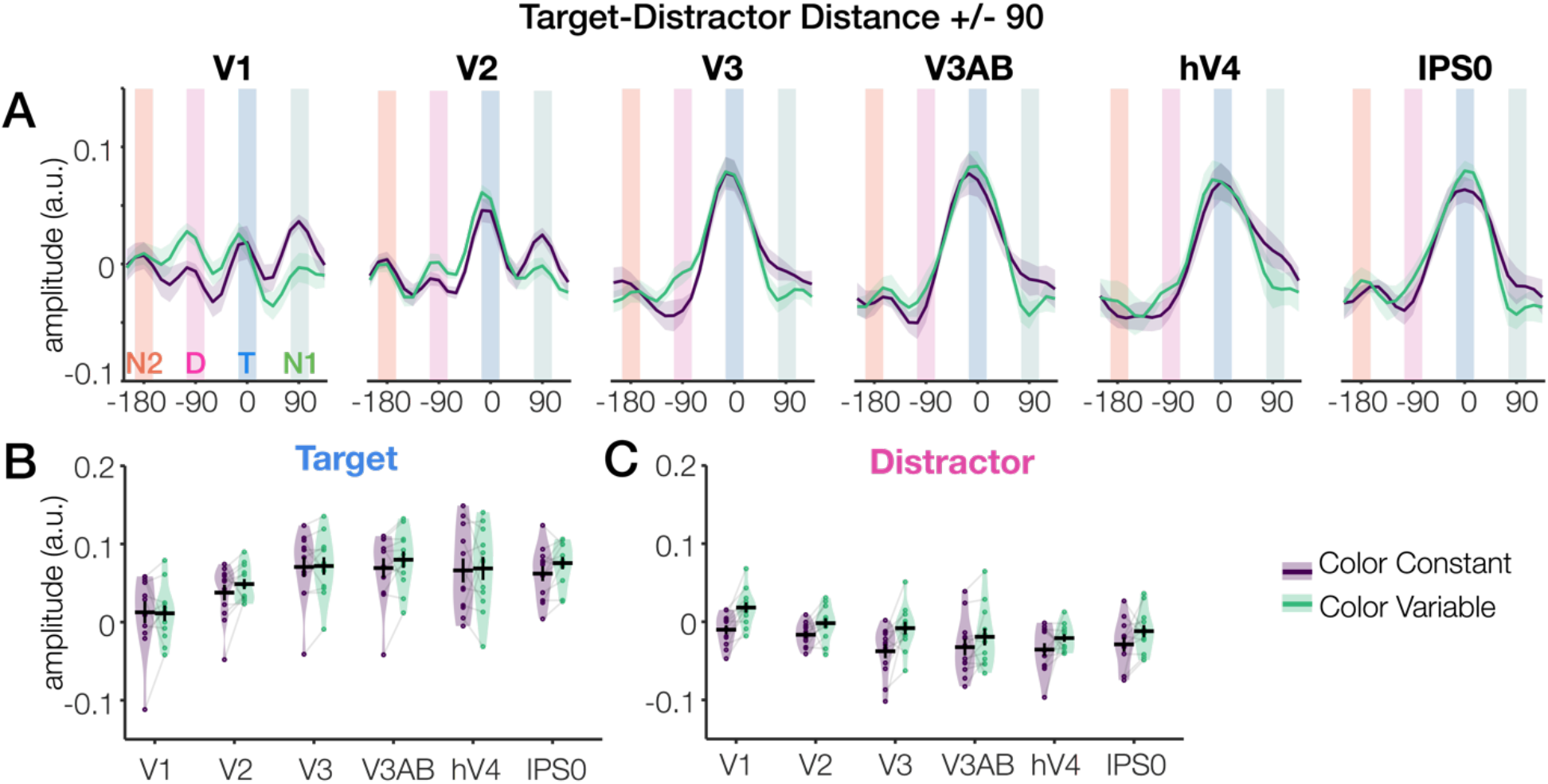
Dissociable effects of stimulus history on target enhancement and distractor suppression. (A) Model responses for individual ROIs as a function of task condition (Arrays with target-distractor distance +/-90). Purple and green lines (Shaded error bars = 1 SEM) show the output of the inverted encoding model in the color constant and color variable conditions, respectively. Background panels at -180°, -90°, 0° and +90° show the positions of the 4 search array items (blue = target (T), pink = distractor (D), green = non-target 1(N1), orange = non-target 2 (N2). Target enhancement can be seen as the greater height at position 0: The IEM peak at the blue bar is higher than the IEM peak at the orange, pink, and green bars. History-driven distractor suppression can be seen as the lower height at position -90 for the color constant vs. color variable conditions: The IEM peak at the pink bar is higher for the green line than for the purple line. (B) Target amplitude as a function of ROI and task condition. There was no effect of task condition on target amplitude, but a significant increase in target amplitude across ROIs. Violin plot shading shows range and distribution of the data; dots represent single subjects; black error bars indicate ±1 SEM. (C) Distractor amplitude as a function of ROI and task condition. There was a significant effect of task condition on distractor amplitude, and this history-driven effect did not interact with ROI.

### Analysis of search array locations in V1, V2, V3, V3AB, hV4, and IPS0

Given that we can assess differential responses associated with each of the 4 items in the search array (Figure 4), we next tested whether goal- and history-driven modulations were differentially represented across the visual stream by performing an analysis of history-driven effects on target and distractor processing across visual ROIs (Figure 5). These six ROIs (V1, V2, V3, V3AB, hV4 and IPS0) were chosen for each participant having at least 90 spatially selective voxels as determined by the localizer data. Here, we focus on history-driven effects on target processing and distractor processing for the arrays where behavioral and neural distractor competition effects were greatest (target-distractor separation +/-90°, see Figure 2E,F). Full ANOVA results and additional plots are shown for individuals ROIs with both array 90° and 180° configurations in extended data Figures 5-4 and 5-5.

We found evidence for within-display target enhancement (i.e., enhancement of the target over other positions), but we did not find evidence for history-driven modulations of target enhancement. Overall target enhancement was significant in all ROIs (all *p*’s < .001) except for V1 (*p*’s > .12), and target enhancement significantly increased across ROIs (*p* < .001) as shown in Figure 5A-B. There was, however, no meaningful effect of history on target amplitude as revealed by a repeated-measures ANOVA testing the main effect of history and the interaction between history and ROI on target processing (*p* = .35, η^2^_p_ = .08; *p* = .64, η^2^_p_ = .04 for main effect and interaction respectively). This pattern was the same whether we used raw amplitude values or we used values from the GLM (no effect of history, *p* = .28, no interaction of history and ROI, *p* = .51).

In contrast, history had a significant effect on distractor amplitude such that distractor amplitudes were significantly attenuated in the color constant condition relative to the color variable condition. A repeated-measures ANOVA revealed a main effect of history (*p* = .007, η^2^_p_ = .50) and no interaction between history and ROI (*p* = .44, η^2^_p_ = .08), indicating that the effect of history on distractor processing was similar throughout the examined ROIs. Though the ANOVA suggests that history effects were of a similar magnitude across all examined ROIs, a post-hoc simple main effects analysis showed that the effect was individually significant only in V1 (*p* < .001) and V3 (*p* < .01). This general pattern was the same whether we used raw amplitude values or else used values from the GLM approach (main effect of history, *p* = .01, η^2^_p_ = .47, no interaction of history and ROI, *p* = .87, η^2^_p_ = .03).

Finally, we examined changes in non-target responses. For “non-target 1” (the item neighboring the target on the side opposite the distractor), there was an overall history related modulation (color constant > color variable, *p* = .016, η^2^_p_ = .42) that did not interact with ROI (*p* = .76, η^2^_p_ = .03). Similar general effects on non-target processing have been observed recently (Won et al., 2020) and may reflect a bias of attention away from the distractor such that attention may ‘overshoot’ the target because of the reduction in signal at the distractor location. The effect of history on “non-target 1” responses likewise was similar though of borderline significance in the GLM analysis (color constant > color variable, *p* = .049, η^2^_p_ = .31). We found no effect of history on the other non-target (“non-target 2”) which occupied the spatial position 180 degrees from the target item (*p* >= .61).

Finally, additional analyses on larger, aggregate ROIs (V1-V3, IPS0-3) yield convergent results and also demonstrate how distractor suppression effects were absent for arrays where the target and distractor did not compete with each other (target-distractor separation +/-180°), consistent with our separate analysis of each ROI (Figure 6) and prior behavioral and neural findings (Turatto and Galfano, 2001; Wang and Theeuwes, 2018a; Failing et al., 2019b; Won et al., 2020) (extended data Figures 5-1, 5-2, and 5-3).

**Figure 6.**
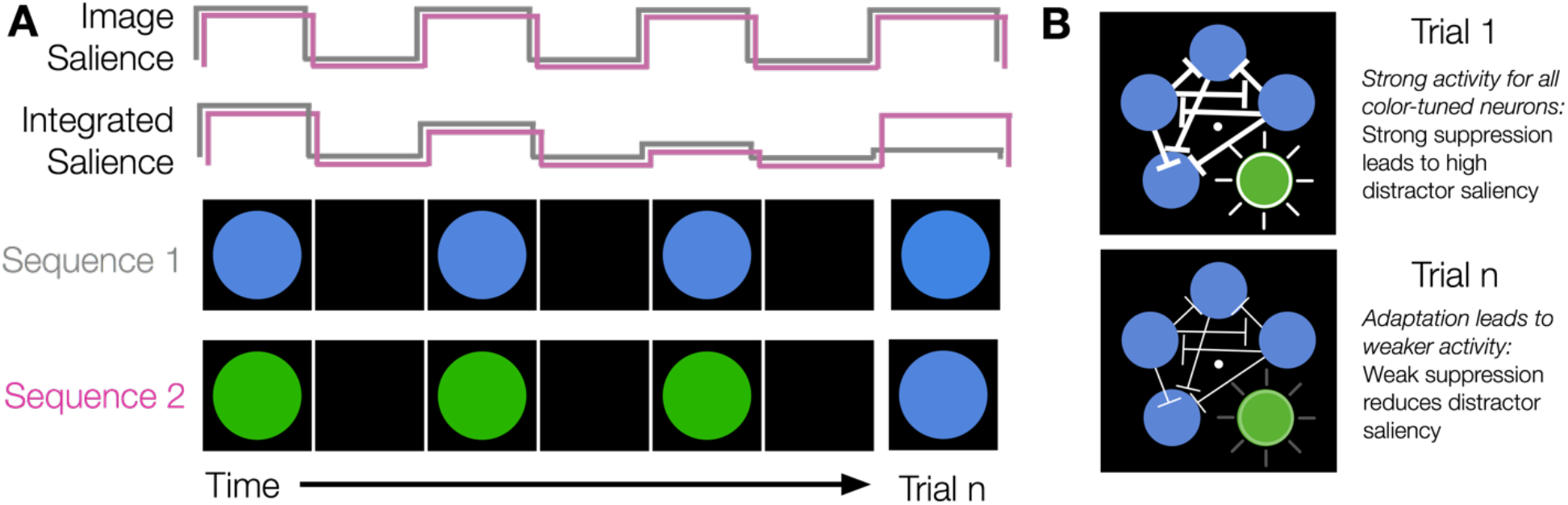
Simplified cartoon illustration of local-image versus temporal-integration salience for a simple image with one feature and location. (A) In 2-D salience computations, stimulus-driven stimulus drive is determined locally within a given image without respect to prior images. Sequence 1 is 4 different trials, and on each trial the same stimulus is shown (Blue-Blue-Blue-Blue). Sequence 2 is 4 different trials, but the final trial is a different color from the preceding trials (Green-Green-Green-Blue). The final trial (Blue) is physically identical for the two sequences. So, the final stimuli (trial n in each sequence) have identical 2-D salience. Assuming that we chose equiluminant green and blue values, then each “frame” in the sequence likewise has approximately the same image-computable salience, as shown by the uniform-sized square pulses in the cartoon. Alternatively, stimulus-driven salience maps may better be conceived of as reflecting a temporally-integrated 3-D salience map, as early sensory neurons adapt to ongoing stimulus features. In Sequence 1 (Blue-Blue-Blue-Blue), the activity of neurons that are maximally responsive to blue wanes due to adaptation. In Sequence 2 the activity of neurons maximally responsive to green wanes over the first 3 trials, but the final stimulus elicits a robust response from the non-adapted blue-preferring neurons. Thus, temporally-integrated salience for the trial *n* in each sequence differs across the two sequences even though the stimuli are physically identical. (B) Most studies of predictive coding and adaptation consider changes to neural activity for a single item. Here, we illustrate how adaptation can have consequences for stimulus-driven saliency that arises from inter-item competition within multi-item arrays (e.g., Itti and Koch, 2000). Top: For the first presentation of the array, all neurons respond strongly, leading to classic inter-item competition effects that yield high distractor saliency. Bottom: With repeated presentations and adaptation, overall activity and inter-item competition is weakened, yielding a relative attenuation of the distractor.

## Discussion

To find what we are looking for, we must integrate information about stimulus relevance, salience, and history. While the impact of stimulus relevance and salience on topographically organized population codes have been thoroughly investigated, stimulus history is not thought to be a wholly goal-driven or stimulus-driven process. Rather, history effects may depend on interactions between the current stimulus drive (‘bottom-up’ factor) and the current internal state of the visual system (‘top-down’ factor). To address this ambiguity and to better understand how history impacts visual processing, we tested whether history-driven changes to attentional priority operate in a manner akin to canonically goal-driven and/or to stimulus-driven signatures of priority. To do so, we estimated population-level neural responses evoked by 4-item search arrays across retinotopically-defined areas of occipital and parietal cortex. We found that stimulus history did not modulate the specificity of goal-driven target templates, as goal-driven target enhancement was unaffected by stimulus history. Instead, we found that stimulus history attenuated responses related to distractors throughout the visual hierarchy. These results suggest that stimulus history may influence visual search performance via local competitive interactions within early sensory cortex (i.e., V1), consistent with the V1 salience map hypothesis. Further, we argue that these early competitive interactions cannot be explained by goal-driven predictive coding models.

### Proposed model: Adaptation alters a stimulus-driven salience map in V1

Models of image-computable salience propose that local image statistics determine competitive interactions that give rise to 2D spatial salience maps within V1 (Li, 2002; Zhang et al., 2012) or after integrating feature maps at a later stage of processing (e.g., Treisman and Gelade, 1980; Wolfe, 1994; Itti and Koch, 2000; Carmi and Itti, 2006), and these models do not typically account for the long-term effects of stimulus history. Note, many existing saliency models do account for *short -term* changes to stimuli by coding for dynamic image factors such as motion velocity and flicker (i.e., luminance onset or offset) across movie frames (∼30 ms per frame), (Itti and Koch, 2000; Carmi and Itti, 2006). However, these dynamic feature maps cannot explain history effects that build up over the course of many trials and persist across blank inter-trial intervals. Rather, an additional mechanism is needed to integrate stimulus information over a longer duration. Recent work suggests that neural adaptation – which is linked to the history of prior stimuli – in a subset of tuned neurons may alter stimulus-driven competitive dynamics (e.g., divisive normalization, Carandini and Heeger, 2012) within early visual cortex (Solomon and Kohn, 2014). Thus, to accommodate our observation of history-driven distractor suppression within existing saliency models, we propose that stimulus-driven evoked responses in V1 may be integrated over a longer, multi-trial duration (as opposed to just within a single image; Figure 6) (Karni and Sagi, 1991; Schwartz et al., 2002; Jehee et al., 2012). In the context of models of visual search, this might be comprised of a series of 2D spatial maps that together form a temporally integrated 3D salience map (i.e., salience is computed based on current and prior physical stimulus properties). Consistent with the notion of a 3D salience map, recent behavioral and neural evidence suggests a role for priming and habituation in visual search behaviors (Geyer et al., 2006; Feldmann-Wüstefeld and Schubö, 2016; Turatto and Pascucci, 2016; Turatto et al., 2018; Won and Geng, 2020, also see: Reavis et al., 2016), even when the adapting stimuli are task-irrelevant.

Consistent with a temporally-integrated V1 saliency account of history-driven distractor suppression, we observed history-driven modulations only with sufficient competition (i.e., targets and distractors were closer together) and we observed robust history-driven modulations in V1 in the absence of goal-driven modulations. In line with our findings, prior behavioral work has shown that incidental repetitions of distractor, but not target, features and locations modulate search performance (Geyer et al., 2006; Failing et al., 2019b). Likewise, prior work has shown a rapid suppression of distractor-evoked neural responses (Hickey et al., 2009; Zhang and Luck, 2009; Sawaki and Luck, 2010; Gaspar and McDonald, 2014; Moher et al., 2014; Gaspelin et al., 2015, 2017) and that the likelihood of distraction results in anticipatory changes to distractor, but not target, locations (Serences et al., 2004; Heuer and Schubö, 2019; Won et al., 2020). However, the proposed temporally-integrated salience account does not capture all history-driven effects. In our task, the repeated distractor features were purely visual in nature, and thus history effects might be mediated entirely via local circuit dynamics (i.e., the adaptation account described above). In contrast, other studies have examined history-driven effects for more abstract features like reward (Mazer and Gallant, 2003; Serences, 2008; Saproo and Serences, 2010; Stanisor et al., 2013; Chelazzi et al., 2014; Hickey and Peelen, 2015; MacLean and Giesbrecht, 2015; Itthipuripat et al., 2019; Kim and Anderson, 2019) (but also see: Maunsell, 2004; Anderson and Kim, 2019), which may require an intermediary pathway such as the medial-temporal lobe (Theeuwes, 2019) or dopaminergic midbrain structures (Hickey and Peelen, 2015, 2017).

### Implications for predictive coding theories of visual processing

Much of the debate about history-driven changes to visual search has been separated from the predictive coding literature, but these two ideas are highly intertwined. Predictive coding theories propose that incoming visual information is compared to expected visual information at later stages of processing (Rao and Ballard, 1999; Friston, 2005; Summerfield and de Lange, 2014; Spratling, 2017). Efficient, predictive coding is achieved via an iterative updating process whereby error units detect deviations from what is expected and inhibit expected information in the prediction units at an earlier stage of processing. Here, we consider whether an expectation-driven predictive coding account, whereby top-down expectations about the upcoming stimulus influence neural processing, could likewise explain the pattern of results that we have observed. We note that the term “predictive coding” has been used in a wide variety of ways in the literature (Spratling, 2017), some of which are entirely stimulus-driven (e.g., within the retina; Srinivasan et al., 1982). Here, we are concerned with considering versions of predictive coding whereby top-down expectations influence stimulus processing.

A fundamental tension has long been noted in the literature: Predictive coding models primarily explain how expected information becomes *attenuated*, and thus have difficulty explaining signal enhancement related to attention (e.g., Luck et al., 1997; Hupé et al., 1998; Kastner and Ungerleider, 2000). Many predictive coding models implement error-driven feedback as inhibitory signaling from the next adjacent visual area (e.g., V3 to V2). To explain top-down attentional prioritization effects, predictive coding models must be modified, as has been done in the Predictive Coding/ Biased Competition (PC/BC) model (Spratling, 2008). In this model, an additional top-down attention component is added, and the error and prediction units are shifted such that error-driven feedback is excitatory. These changes to the model allow for biased competition effects to arise within a predictive coding framework.

Although the PC/BC model variants can predict biased competition effects in attention, such models critically predict that target enhancement and distractor suppression effects will be yoked, as both effects arise from feedback from the next higher level of visual processing. Thus, for existing predictive coding models to explain our results, we should have observed that the emergence of history-driven distractor suppression paralleled top-down target enhancement. In contrast, we found diverging target enhancement and history-driven distractor suppression effects: whereas target enhancement was absent in V1 and increased across the visual stream, history-driven distractor suppression emerged in V1. Thus, we propose that history-driven distractor suppression is best explained by ‘bottom-up’ inter-trial priming arising from adaptation within V1 (Westerberg et al., 2019).

Furthermore, we argue that it is important to differentiate between “bottom-up” and “top-down” expectational effects, analogous to recent arguments that it is critical to differentiate between potential confounds of attention and expectation (Summerfield and de Lange, 2014; Rungratsameetaweemana and Serences, 2019). We define ‘top-down’ expectations as those that can be updated flexibly and on a rapid time scale (e.g., over the course of a few trials). In contrast, we define ‘bottom-up’ expectations as those that are ingrained over a very long time-scale and tied to particular stimuli. For example, early ideas about predictive coding emerged from studies of the retina: By exploiting long-term ‘expectations’ that naturalistic stimuli are correlated in space and time, coding within the retina can be highly efficient (Srinivasan et al., 1982; Rao and Ballard, 1999).

Making a distinction between ‘bottom-up’ and top-down’ expectations can explain prior results that run counter to some predictive coding models. Specifically, Maljkovic and Nakayama’s (1994) priming of pop-out experiments demonstrated that RT costs are incurred by switching stimuli even when the stimulus switch is expected. When a stimulus is predictable and repeated (e.g., 0% probability of a color switch), participants are faster than when a stimulus is unpredictable and switches color (e.g., 50% probability of a color switch. If expectations can attenuate the cost of switching colors, then participants should likewise be faster in a predictable, 100% switch condition than in the unpredictable 50% switch condition. In contrast to this prediction, Maljkovic & Nakayama found that participants were *slower* in the 100% switch condition: Participants apparently were unable to use their expectations to overcome bottom-up stimulus-driven priming effects.

### Goal-driven attention effects

In addition to implicating early visual cortex in representing history-driven task factors during visual search, we also replicated prior findings that the locations of attended items (here, search targets) are prioritized relative to other item locations in both visual and parietal cortex (Saproo and Serences, 2010; Sprague and Serences, 2013; Sprague et al., 2018b). These target-related modulations are consistent with the broad involvement of visually-responsive regions in representing goal-driven priority during visual search (Mazer and Gallant, 2003; Ogawa and Komatsu, 2006). For example, recent studies manipulated the salience (contrast) and relevance (attended or unattended) of items and found that salience and relevance were both represented, to varying degrees, across the visual hierarchy (Poltoratski et al., 2017; Sprague et al., 2018b). Notably, however, here we found that target prioritization was absent in V1, whereas prior work has found robust effects of attention in V1 (Motter, 1993; Kastner, 1998; Tootell et al., 1998; Gandhi et al., 1999; Kastner et al., 1999; Somers et al., 1999; Serences and Yantis, 2007; Saproo and Serences, 2010; Sprague and Serences, 2013). This difference may reflect task differences — much prior work found attention-related gains in V1 when spatial attention was cued in advance or a single target was shown, whereas visual search arrays provide visual drive at many competing locations and spatial attention is deployed only after array onset. As such, further work may be needed to unconfound history effects and attention effects in the study of spatial attention, as much early work on univariate attention effects has employed blocked designs where the same location is attended for many trials in a row (Kastner, 1998; Tootell et al., 1998; Gandhi et al., 1999; Kastner et al., 1999; Somers et al., 1999).

### Future directions

Although our work suggests that stimulus history modulates representations of distractor but not target processing in visual cortex, there are some potential limitations to the current design that suggest avenues for future work. First, because we measured only location, we could not directly measure suppression of the distractor color (Failing et al., 2019a). However, as the spatial position of the distractor was completely unpredictable, our results do strongly imply that the distractor color was suppressed. Likewise, most theories of visual search hypothesize that space is the critical binding medium through which feature and goal maps are integrated (Treisman and Gelade, 1980; Wolfe, 1994; Itti and Koch, 2000), and recent work suggests that location is spontaneously encoded even when only non-spatial features such as color are task-relevant (Foster et al., 2017a). Second, it is possible that history may modulate both distractor- and target-processing in other circumstances not tested here. That is, perhaps the target template ‘diamond’ in our task was sufficiently useful such that adding feature information to this template (e.g., ‘red diamond’ rather than ‘diamond’) did not confer a behavioral advantage (but see: Maljkovic and Nakayama, 1994). Finally, the time-course of MRI (sampling every 800 ms) is slower than shifts of spatial attention to the search target (< 500 ms) (Foster et al., 2017b). Although the history-driven effects that we observed in visual cortex are consistent with the rapid distractor suppression effects observed in EEG (Sawaki and Luck, 2010; Gaspar and McDonald, 2014), we cannot definitively say on the basis of these data that the observed history-driven effects occurred rapidly and directly within visual cortex versus via recurrent feedback from later visual areas. Nonetheless, the present work is consistent with and provides critical initial evidence for such a model.

## Acknowledgements

We thank Rosanne Rademaker for scanning assistance and for sharing custom analysis code. We thank Nicole Rangan and Matteo d’Amico for assistance with behavioral data collection, and we thank Ed Awh for helpful comments on earlier drafts of the manuscript.

