## Extended Data for "History modulates early sensory processing of salient distractors"

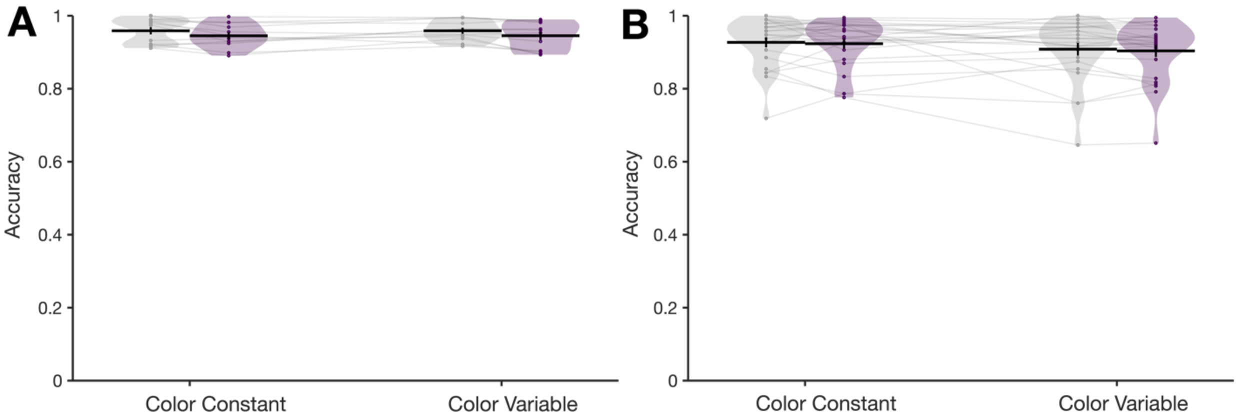


**Figure 2-1. Accuracy measure for the behavior data. (A)** Accuracy by task condition in the fMRI experiment. **(B)** Accuracy by task condition in the behavioral experiment. We performed a repeated measures ANOVA with factors Task Condition (color constant, color variable) and Distractor (present, absent). In the MRI sample (Experiment 1A), we found no main effect of task, $F(1,11) = <.001$, $p = .99$, $\eta^2_p = <.001$. We found a main effect of distractor that was inconsistent with a speed-accuracy tradeoff; participants were slightly less accurate on distractor present trials (M = 94.5%, SE = 0.9%) than on distractor present trials (M = 95.9%, SE = .9%), $F(1,11) = 14.31$, $p = .003$, $\eta^2_p = .57$. We found no interaction between task and distractor, $F(1,11) = < .001$, $p = .98$, , $\eta^2_p = <.001$. In the behavioral sample (Experiment 1B), we likewise found no effect of task, $F(1,23) = = 2.12$, $p = .16$, $\eta^2_p = .08$ and no interaction of task and distractor, $F(1,23) = 0.01$, $p = .92$, $\eta^2_p < .001$. In this sample, however, we failed to find a main effect of distractor, $F(1,23) = 0.93$, $p = .35$, $\eta^2_p = .04$, with mean accuracies of 91.8% (SE = 1.4%) and 91.4% (SE = 1.4%) for distractor absent and present trials, respectively.

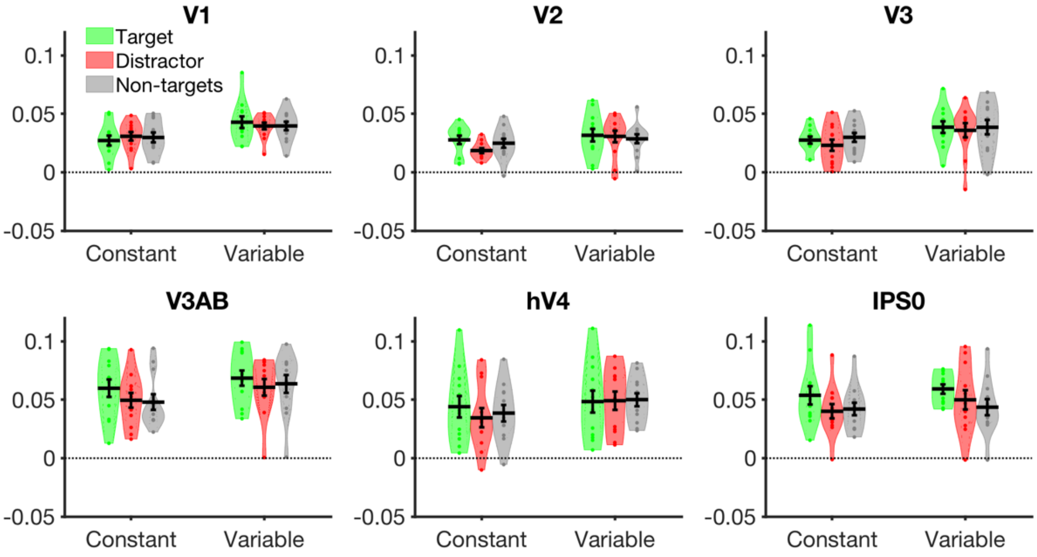


**Figure 3-1. Univariate response in voxels selective to each item position, as a function of item type (target, distractor, non-target) and ROI.** We identified voxels that were selective to one quadrant over the others using the independent mapping task (e.g., voxels that showed greater activity when the wedge stimulus was presented in quadrant 1 versus the other quadrants). We then created event-locked time courses for each trial (baselined to the first TR), and we averaged the univariate activity for the voxels for each item position (TRs 4-10, same as in the classification analysis). A repeated-measures ANOVA with factors ROI, Item (target, distractor, non-target) and Condition (color constant or color variable) revealed a main effect of ROI (greater activation for later areas, $p < .001$), a main effect of item ($p = .022$) and no main effect of condition ($p = .09$; though, this was numerically in the direction of less overall activity in the constant condition, i.e., univariate repetition suppression). No interactions reached significance. A simple main effects analysis of the item effect revealed that it was driven entirely by IPS0 ($p = .007$); there was no significant effect of item in any other ROI ($p >= .19$).

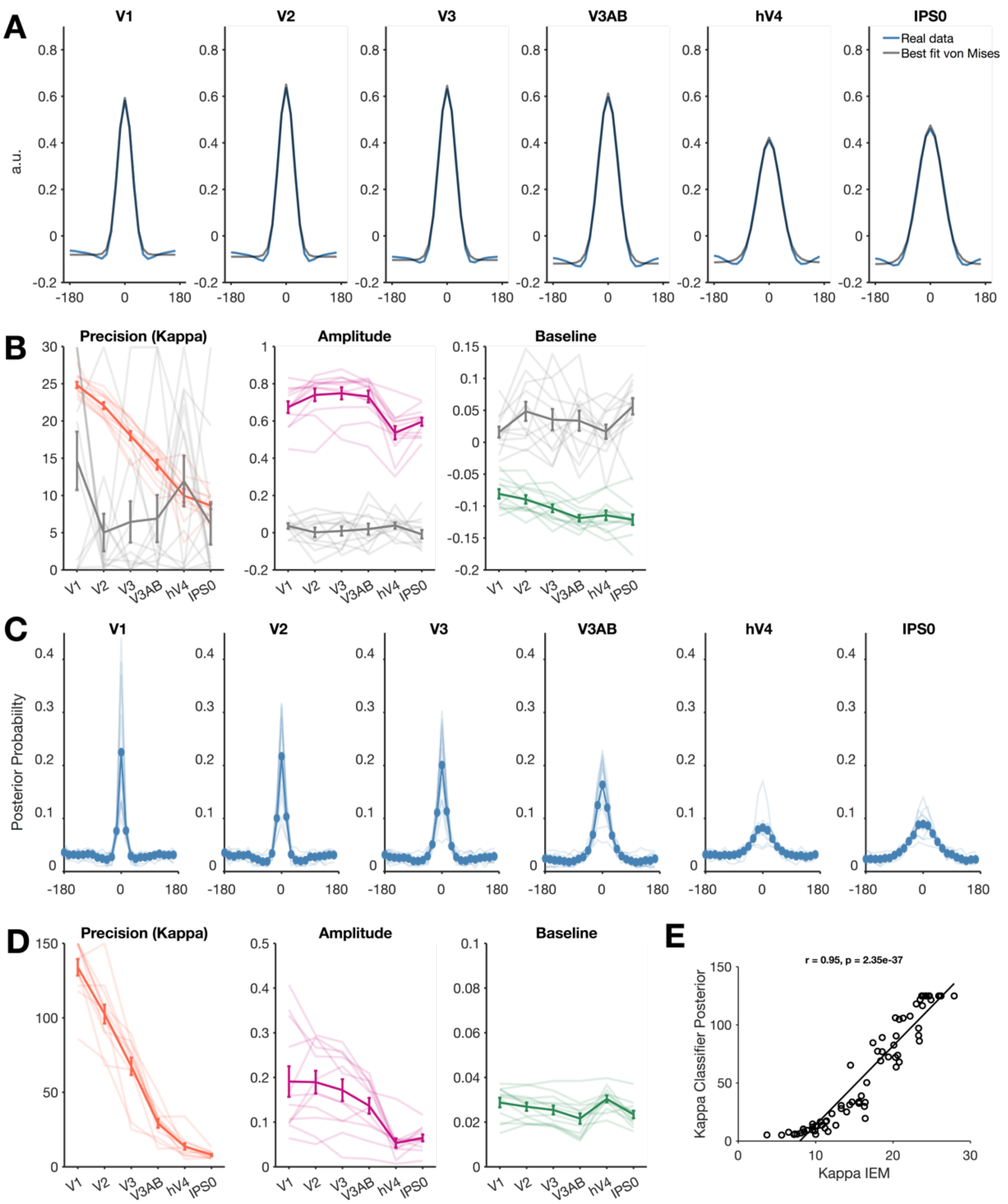


**Figure 3-2. Average best-fitting von Mises distribution, linear classifier posterior probabilities and parameter estimates. (A)** Single-item model estimates based on training and testing within the independent mapping task data. We fit von Mises distributions to inverted encoding model fits, training and testing within the independent mapping task data (parameters shown in Figure 3). Best fitting distributions are depicted in Figure S1. To fit the von Mises to the inverted encoding model estimate for each ROI, we performed a grid search over a range of possible precision values (kappa = 0.001 to 30 in increments of 0.1). For each precision value, we chose the best fit baseline, amplitude, and central tendency (mu) values and calculated root mean square error to quantify fit. **(B)** Plot of best-fitting parameters with additional gray lines to show the best-

fitting parameters to the shuffled data. Consistent with the shuffled data being better fit by a uniform distribution than by a 3-parameter von Mises, the amplitude and baseline parameters held near 0 for the shuffled data whereas the precision parameter was highly unstable. **(C)** Posterior probabilities of each position bin being chosen by a linear classifier (*classify.m, 'diagLinear'* covariance option). **(D)** Best fitting von Mises parameters when fitting a von Mises to the posterior probabilities obtained by the linear classifier. These parameters closely parallel changes to IEM estimates across ROIs, but with a change in scale, as shown in: **(E)** Correlation between the Kappa parameter for a von Mises fit to IEM output versus linear classifier posterior probabilities.

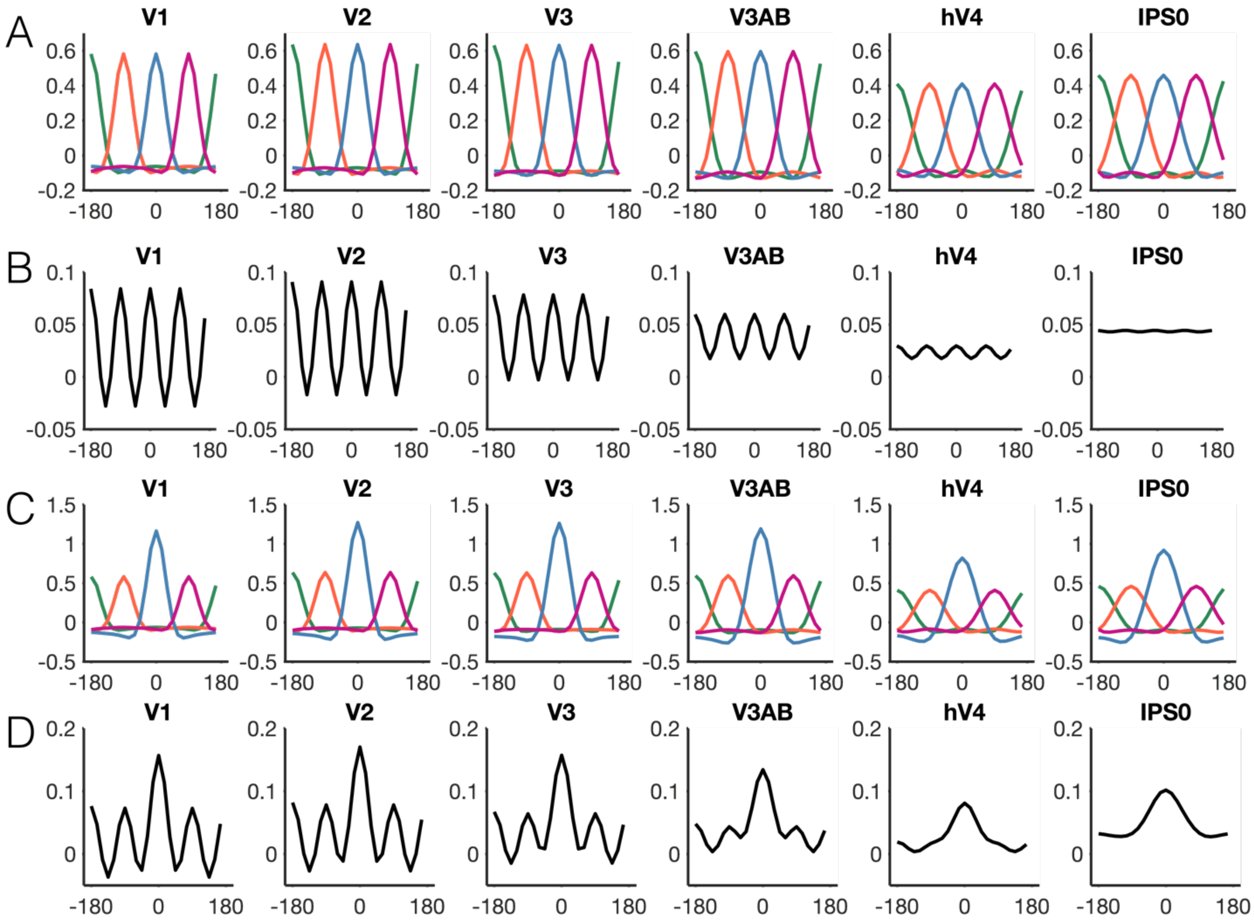


**Figure 4-1. Generating predictions for 4-item model estimates by averaging single-item model estimates, all ROIs.** (A) Average from the independent mapping task plotted at 4 hypothetical item locations. Here, these 4 "items" are represented with equal priority. (B) Hypothetical observed response when measuring a single trial containing the 4 items presented simultaneously. This line is the average of all lines in Panel A. (C-D) The same as panels A and B, but with the item at position 0 assigned a higher "priority" (i.e., gain) than the other three items.

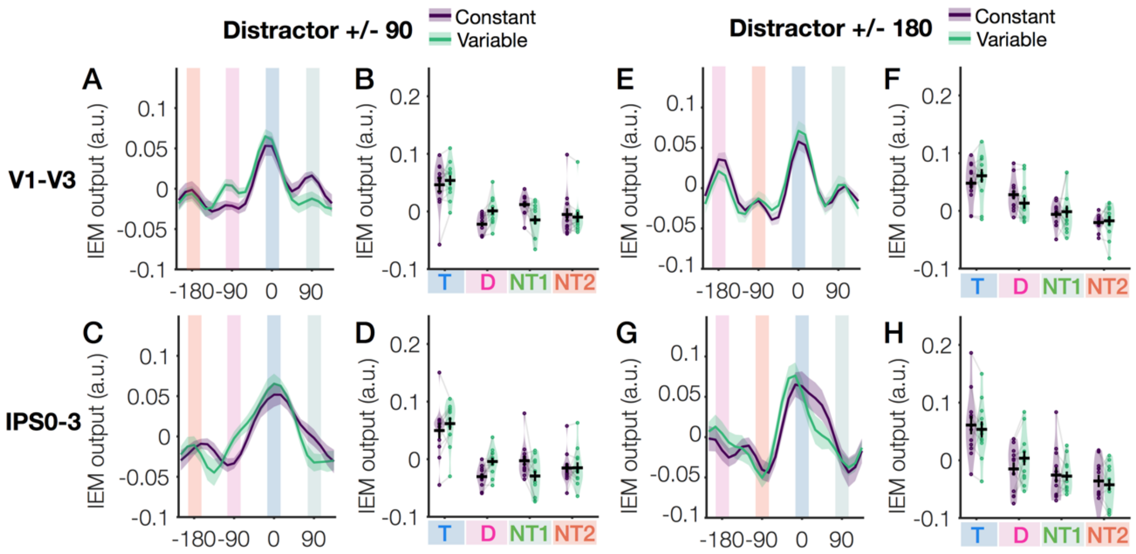


**Figure 5-1. Aggregate ROI analysis reveals target enhancement and distractor suppression in both early visual and parietal cortex.** (A) Model estimates for all channels in early visual cortex for arrays with target-distractor distance of +/- 90°. Lines show model estimates for the color constant and color variable conditions (B) Model estimated at expected item peaks (+/- 15° from peak). Color-coded background panels indicate the positions of the search array items. Symbols indicate uncorrected comparisons between conditions at each item location: *n.s.* $p$ >.10, ~ $p$ < .10, *$p$ < .05, **$p$ < .01, ***$p$ < .001). (C-D) Model estimates for all channels (C) and expected item peaks (D) in parietal cortex for target-distractor distance of +/- 90°. (E-F) Model estimates for all channels (E) and expected item peaks (F) in early visual cortex for arrays with target-distractor distance of +/- 180°. (G-H) Model estimates for all channels (G) and expected item peaks (H) in parietal cortex for target-distractor distance of +/- 180°

An initial ANOVA with factors Array Configuration (target-distractor distance of 90° versus 180°), Condition (color constant or variable) and Item (position of the target, distractor, non-target 1, or non-target 2) interacted with several aspects of the data, complicating interpretation. These interactions include an interaction of Array Configuration and ROI, $F(1,11) = 15.10$, $p = .003$, $\eta^2_p = .58$, Array Configuration and Item, $F(1,11) = 4.25$, $p = .012$, $\eta^2_p = .28$, and a 3-way interaction with our key effect of interest (Array Configuration x Item X Condition), $F(3,33) = 3.15$, $p = .038$, $\eta^2_p = .22$. To more easily interpret our key interaction of interest (Item x Condition), we separately examined each Array Configuration (90° or 180° target-distractor separation).

A repeated measures ANOVA on only the 90° target-distractor separation revealed an effect of history on distractor suppression, but not target enhancement in both visual and parietal cortex (A-D). We found a main effect of Item ($p < .001$) driven by the target relative to all other items ($p_{holm} < 1x10^{-5}$), and we found an interaction of Item x Condition, $F(1.99, 21.89) = 9.72$, $p < .001$, $\eta^2_p = .47$. We used a simple main effects analysis (effect factor: Condition, moderator factor: Item), to demonstrate that the Item x Condition interaction was driven by changes to the distractor position, $p < .001$, and to the non-target position opposite the target, $p = .009$ (but not to the target or second non-target, $p >= .14$). The trial-wise physical displays were identical for these two conditions. Thus, model estimates of a physically identical distractor position were modulated by trial history (color constant vs. variable), consistent with suppression of the distractor position in the color constant condition. There was, however, no effect of trial history on target amplitude. There was no three-way interaction of ROI x Item x Condition ($p$ = .98), indicating that this effect was consistent across the aggregate visual and parietal ROIs. ANOVAs within each ROI revealed that the target prioritization effect was present ($p$ < .001). However, the observed distractor suppression effect (Item x Condition interaction) was only found in visual cortex ($p < 1\text{x}10^{-5}$) and absent in parietal cortex ($p$ = .69) when compared to the distractor absent baseline. The Item x Condition interaction in visual cortex (A-B) was driven by a lower distractor value in the color constant condition compared to the distractor absent condition (simple main effects; $p$ = .002), as well as by lower target ($p$ = .002) and higher non-target 1($p$ < .001) values.

Consistent with behavior, an analysis of the 180° target-distractor separation arrays suggest that distractor suppression is needed only when target-distractor competition is sufficiently strong. For the 180° target-distractor separation arrays we found evidence of target enhancement, but no evidence of history-driven distractor suppression (E-H). We again found a main effect of Item, $p < 1\text{x}10^{-6}$) driven primarily by the target being higher than all other items ($p_{holm} < 1\text{x}10^{-5}$), and there was also some evidence that the distractor was greater than at least one non-target location (D vs. NT1 $p_{holm}$ = .012; D vs. NT2 $p_{holm}$ = .13). Unlike the 90° arrays, however, we found no interaction of Condition with the Item effect, $F(3,33)$ = <.001, $p$ = .99, $\eta^2_p$ = .003, and no other effect or interaction with Condition ($p$ > .05). Together, this suggests that the three-way interaction of Array Configuration x Item x Condition in the initial ANOVA was driven by a strong history-driven distractor suppression effect for the 90° array configuration but not for the 180° array configuration. ANOVAs within each ROI revealed that the target prioritization effect was present in both individual ROIs ($p$ < .001). However, the observed distractor enhancement effect (Item x Condition interaction) was only found in parietal cortex ($p$ = .001) and absent in visual cortex ($p$ = .27) when compared to the distractor absent baseline. In parietal cortex (G-H), the Item x Condition interaction was driven by a higher distractor value in the color variable condition compared to the distractor absent condition (simple effects analysis; $p$ = .002) and a lower +90 non-target value ($p$ = .035).

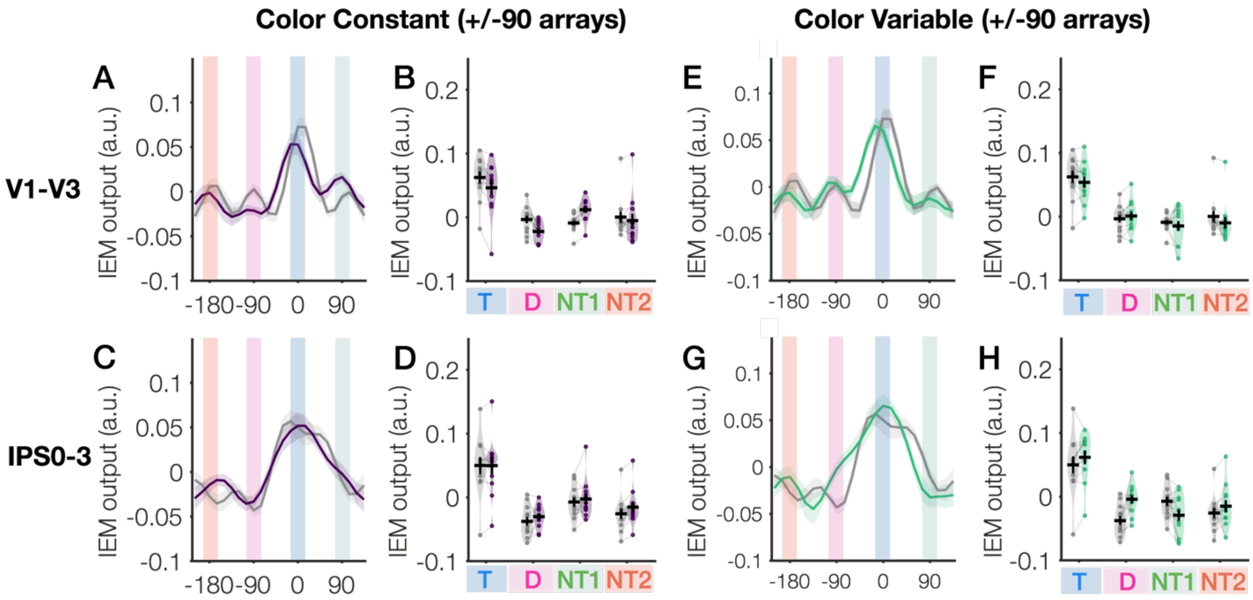


**Figure 5-2. Comparison of each distractor-present task condition to a distractor-absent baseline (+/- 90 degree arrays).** (A-B) Model estimates for all channels (A) and expected item peaks (B) in early visual cortex (V1-V3), comparing the color constant condition (distractor present, target-distractor distance +/- 90) to a target absent baseline. (C-D) Model estimates for all channels (C) and expected item peaks (D) in parietal cortex, comparing the color constant condition (distractor present, target-distractor distance +/-180) to a target absent baseline. (E-H) Plots from A-D for the color variable condition.

Comparing distractor present displays as a function of task condition (Figure 5-1) is most ideal because it ensures that single-trial physical display differences are controlled. However, we also wanted to test that using a distractor absent baseline yields a similar pattern of results. Note, in the distractor absent baseline, there are 3 non-target positions (i.e., there is no true "distractor" position, because all non-targets are physically identical). For the purposes of this analysis, we labelled one of these non-target positions as the "distractor" for the distractor absent baseline. We used the non-target position at the same spatial position relative to the target as in the corresponding distractor present trials (e.g., 90 degrees from the target).

We first considered the color constant condition relative to the distractor absent baseline (A-D). Based on prior analyses and the behavior, we expected to observe distractor suppression relative to a distractor absent baseline. As expected, we again found evidence for overall target enhancement (main effect of Item, $p < .001$; Target greater than all other locations, $p_{holm} < .001$). We likewise again found a pattern consistent with suppression of the distractor location, as indicated by an interaction of Item and Condition ($p = .008$). Here, however, we also found a main effect of ROI ($p = .008$), an interaction of condition and ROI ($p < .001$) and a 3-way interaction of ROI, condition, and item ($p = .006$).

We next considered the color variable condition (E-H). Based on the behavior, we expected to observe distractor *enhancement* in this condition relative to a distractor absent baseline (i.e., capture). We again found overall evidence for target enhancement (main effect of Item, $p < .001$; Target greater than all other locations, $p_{holm} < .001$). We also observed model estimates consistent with distractor enhancement in the color variable condition relative to the distractor absent condition. There was an interaction of Item and Condition ($p$ = .004), and this interaction was driven by higher values for the distractor in the color variable condition versus the distractor absent baseline (simple effects analysis; $p$ = .001). We again found a main effect of ROI ($p$ = .008), an interaction of condition and ROI ($p$ < .001) and a 3-way interaction of ROI, condition, and item ($p$ = .008).

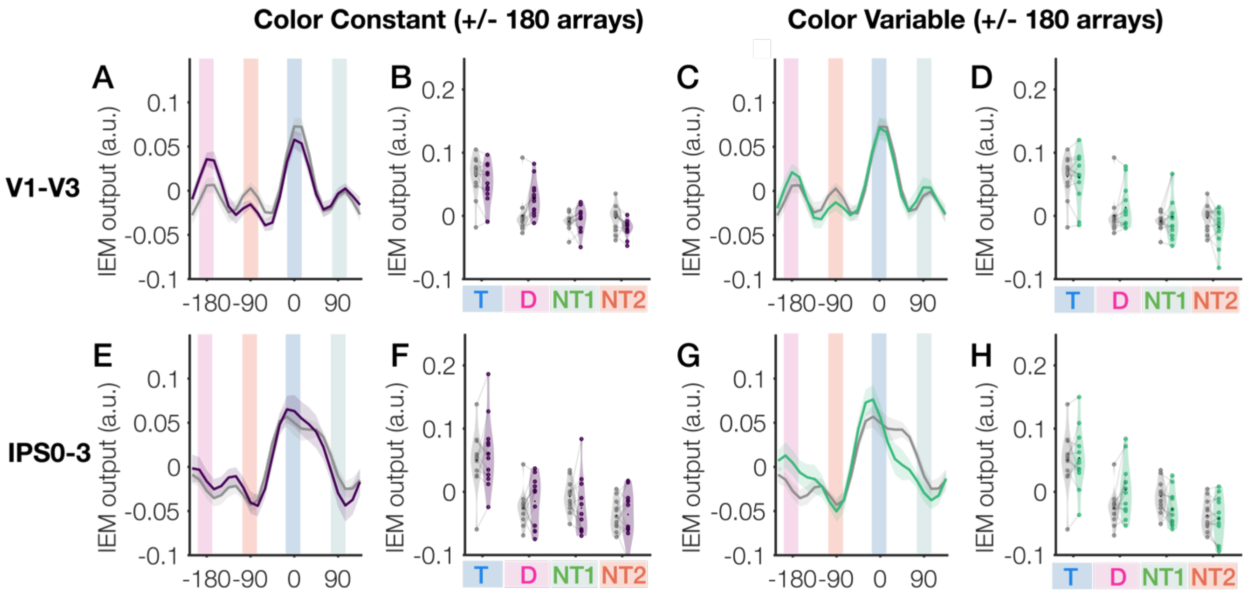


**Figure S5-3. Comparison of each distractor-present task condition to a distractor-absent baseline (+/- 180 degree arrays).** (A-B) Model estimates for all channels (A) and expected item peaks (B) in early visual cortex (V1-V3), comparing the color constant condition (distractor present, target-distractor distance +/- 180) to a target absent baseline. (C-D) Model estimates for all channels (C) and expected item peaks (D) in early visual cortex, comparing the color variable condition (distractor present, target-distractor distance +/- 90) to a target absent baseline. (E-H) Plots from A-D for parietal cortex (IPS0-3).

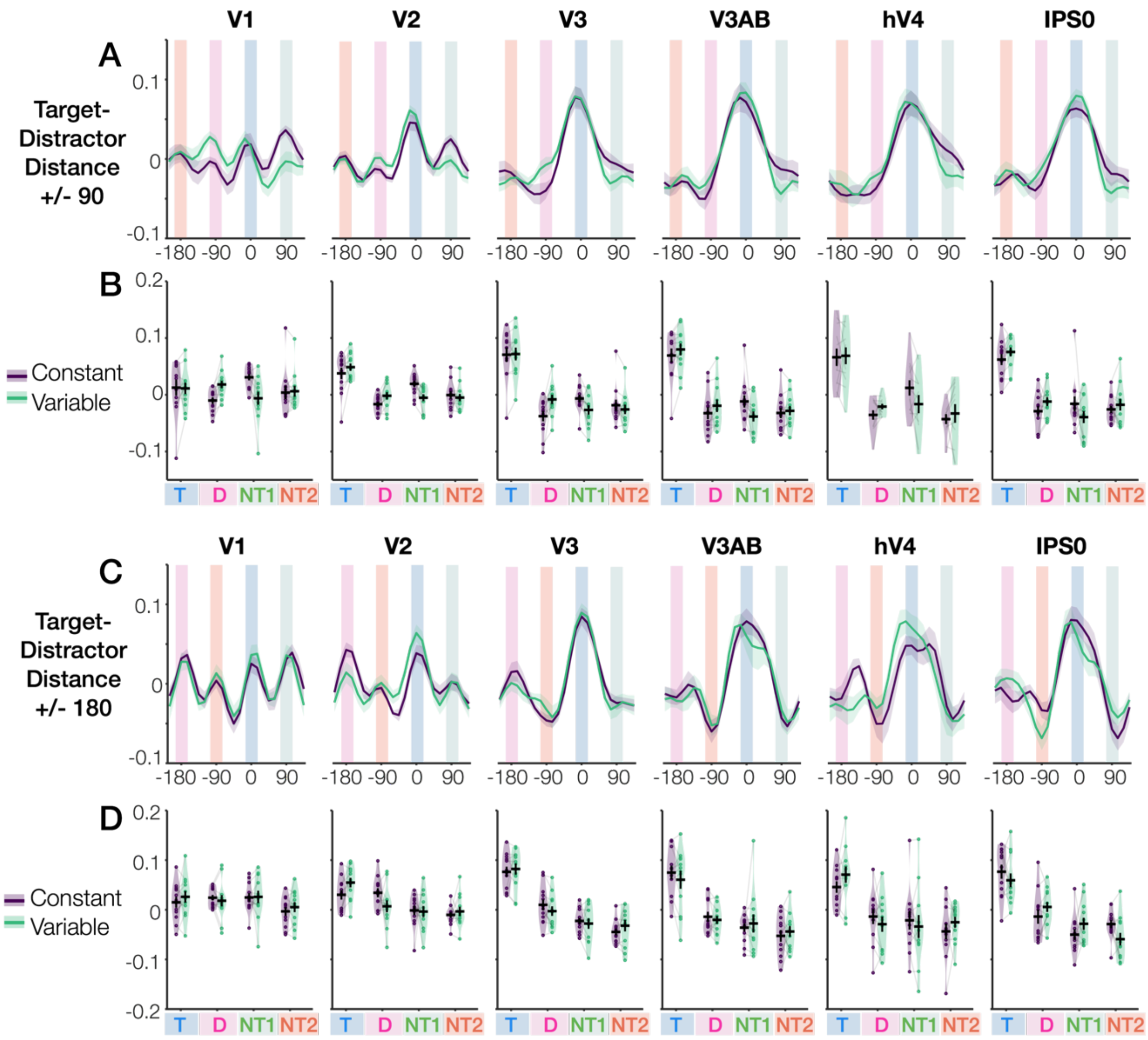


**Figure 5-4. Individual ROI analysis for both array configurations.** (A) Model estimates for all channels in early visual cortex for arrays with target-distractor distance of +/- 90 degrees. Lines show model estimates for the color constant and color variable conditions. Color-coded background panels indicate the positions of the search array items. (B) Model estimated at expected item peaks (+/- 15 degrees from peak). (C-D) Model estimates for all channels (C) and expected item peaks (D) in early visual cortex for arrays with target-distractor distance of +/- 180 degrees. (E-F) Model estimates for all channels (E) and expected item peaks (F) in parietal cortex for target-distractor distance of +/- 90 degrees. (G-H) Model estimates for all channels (G) and expected item peaks (H) in parietal cortex for target-distractor distance of +/- 180 degrees.

We again first performed a repeated measures ANOVA with 4 within-subjects factors: ROI, Array Configuration (90 or 180), Condition (color constant or variable), and Item (target, distractor, non-target +90, non-target +180). We again found a main effect of Item ($p < 1x10^{-5}$) which was driven by target enhancement relative to other items ($p_{holm}$'s $< 1x10^{-6}$). We again, however, observed a 3-way interaction of Array Configuration, Condition, and Item ($p$ = .016), complicating interpretation of history-driven effects. As such, we again conducted separate ANOVA's for each array configuration, each with the factors ROI, Condition, and Item.

Consistent with the aggregate ROI analyses for the 90-degree array configuration (A-B), we again found significant target prioritization (main effect of Item, $p < 1x10^{-8}$) and a significant history-driven modulation of item responses (significant Item x Condition interaction, $p = .002$) that was driven by a suppressed distractor ($p = .007$) and enhanced non-target 1 position ($p = .017$) for the color constant condition relative to the color variable condition. There was no interaction of ROI x Item x Condition ($p = .93$), indicating that this history-driven effect did not significantly vary by ROI. We also found some differences across ROIs, as indicated by a significant main effect of ROI (*p = .003),* an interaction of ROI x Item ($p < 1x10^{-15}$). Upon visual inspection of the data, this main effect and interaction were apparently driven by an increasingly robust target representation from early to late visual ROI's. Follow-up analyses confirmed this impression. Namely, polynomial contrasts for factor ROI (V1,V2,V3,V3AB,hV4,IPS0) revealed a significant linear trend, $p < .001$. A simple main effects analysis (simple effect factor Item; moderator factor ROI) revealed that the main effect of Item was absent in V1 (p = .89) but present in all other ROI's ($p < 1x10^{-5}$). A follow-up ANOVA with area V1 alone confirmed that although there was no target prioritization effect in V1, there were still robust history-driven effects on the distractor as indicated by a significant Item x Condition interaction, $p = .003$, and a significant decrease to the distractor position in the constant versus variable task condition, $p < .001$.

Consistent with the aggregate ROI analyses for the 180-degree array configuration, we found a target enhancement effect ($p < .001$) but no history-driven effects (Condition x Item interaction $p = .60$). We did, however, again observe differences across ROIs, as indicated by a significant main effect of ROI ($p < 1 \times 10^{-7}$), an interaction of ROI x Item ($p < 1x10^{-9}$), and a 3-way interaction of ROI, Item, and Condition ($p = .04$). Polynomial contrasts for factor ROI (V1,V2,V3,V3AB,hV4,IPS0) again revealed a significant linear trend, $p < 1x10^{-9}$. A simple main effects analysis (simple effect factor Item; moderator factor ROI) revealed that the main effect of Item was again absent in V1 (p = .27) but present in all other ROI's ($p <= .001$). A follow-up ANOVA with area V1 alone confirmed that there was no Item x Condition interaction ($p = .83$) for the 180 degree array configuration, consistent with the aggregate ROI analyses.

During each IEM fold, we also estimated "underlying item representation" parameters for each of the 4 items locations (i.e., target, distractor, non-target 1, non-target 2) using a non-negative least squares solution to the general linear model. This allows us to extract the contribution of each of the 4 underlying items to the average observed model estimate in a principled way (confirming that we get the same answer as just comparing the height of 4-item estimate peaks in different conditions). This approach yielded results that were consistent with the main analysis which just used the expected item peak (e.g., model output values at the target peak +/- 15 degrees).

Using the "item strength" values derived from the GLM approach, we again ran a repeated measures ANOVA on the aggregate ROI data (V1-V3, IPS0-3) and using only the 90° target-distractor separation arrays (where we observed the key Item x Condition interaction indicating history-driven modulation. We again found evidence for overall target prioritization, as indicated by a main effect of Item ($p < 1x10^{-5}$) driven by the target relative to all other items ($p_{holm} < 1x10^{-4}$). In addition, we again found an interaction of Item x Condition, $F(3, 33) = 6.60$, $p = .001$, $\eta^2_p = .38$. There was no three-way interaction of ROI x Item x Condition ($p = .84$), indicating that this effect was consistent across the visual and parietal ROIs.

We similarly repeated the analysis for individual ROIs (V1,V2,V3,V3AB,hV4 and IPS0). Using the "item strength" values derived from the GLM approach, we again ran a repeated measures ANOVA using only the 90° target-distractor separation arrays (where we observed the key Item x Condition interaction indicating history-driven modulation. We again found evidence for overall target prioritization, as indicated by a main effect of Item ($p < 1x10^{-6}$) driven by the target relative to all other items ($p_{holm} < 1x10^{-4}$). In addition, we again found an interaction of Item x Condition, $F(3, 33) = 5.50$, $p = .004$, $\eta^2_p = .33$. There was no three-way interaction of ROI x Item x Condition ($p = .79$), indicating that this effect was consistent across the visual and parietal ROIs.

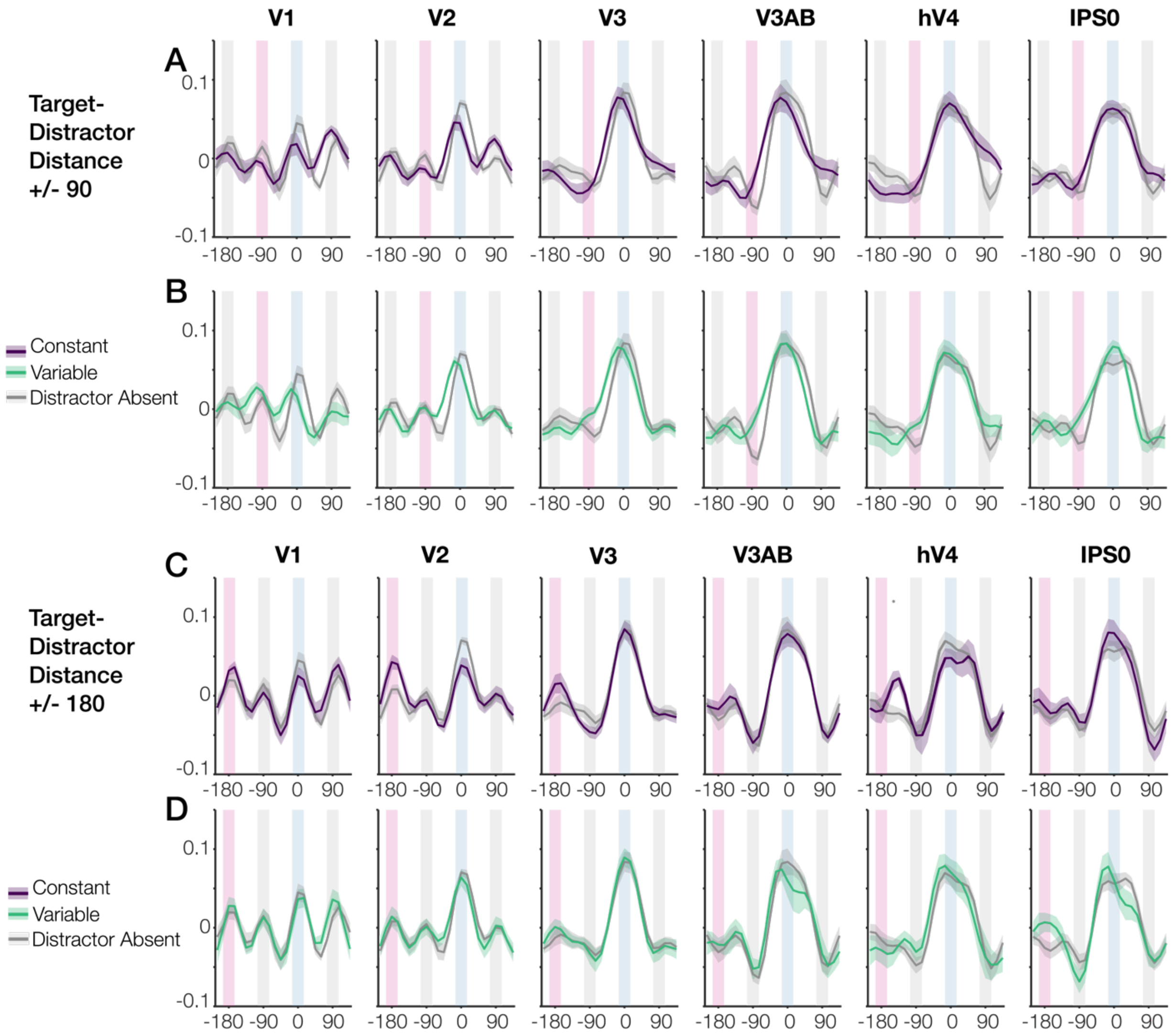


**Figure 5-5. Individual ROI analysis: Each target-present task condition versus a target absent baseline.** (A-B) Comparison of the color constant condition (A) and color variable condition (B) to a target absent baseline when the target-distractor distance was 90 degrees. (C-D**)** Comparison of the color constant condition (C) and color variable condition (D) to a target absent baseline when the target-distractor distance was 180 degrees.
